# AI-discovered protein fragments as generalizable regulators of biomolecular condensates

**DOI:** 10.64898/2026.05.08.723928

**Authors:** Andrew Savinov, Jibin Sadasivan, Kyle J. White, Jack D. Rubien, Gene-Wei Li, Lindsay B. Case

## Abstract

Biomolecular condensates are a major driver of cellular organization; however, we lack a predictable and systematic approach to modulate their underlying multivalent interactions. Here, we demonstrate a generalizable AI-driven method for designing protein fragments to control condensate formation, applying this approach across G3BP1, SARS-CoV-2 nucleocapsid, TDP-43, and focal adhesion kinase (FAK). Computationally screening 2,235 fragments, we selected 18 for experimental investigation, attaining a 50% success rate. Furthermore, predicted fragment binding modes align with their activities, revealing known and novel interactions driving condensate formation. For example, a fragment which suppresses FAK condensates in mammalian cells uncovered an interdomain interaction required for phase separation. Together, our results establish AI-guided protein fragment discovery as a generalizable strategy to dissect and control the molecular interactions that govern biomolecular condensates.

## Introduction

The formation of biomolecular condensates through phase separation is a fundamental principle of cellular organization^1–3^. Condensates concentrate proteins and nucleic acids to form membrane-less compartments with physiological functions including stress responses^4–7^, signal transduction^8^, and immunity^9,10^. Understanding the underlying properties, compositional specificity, and molecular functions of condensates^11–14^ will be critical for understanding cell physiology and rationally treating diseases characterized by aberrant phase separation^15,16^ – including neurodegeneration^17^, cancer^18,19^, and viral infection^20,21^. All of these goals would be well served by a generalizable approach to map and control the interactions driving condensate formation.

Recent work has revealed high-throughput methods to study protein condensates at scale^22–26^, providing one path towards general principles. Such approaches are complementary to the high-resolution biochemistry, cell biology, analytical theory, and biophysical modeling that have been foundational to this field (e.g., refs. 27–31). Protein design methods have also been used to engineer binders targeting intrinsically disordered regions (IDRs) to label condensates and in some cases disrupt them^32^. However, synthetic binders provide insight into only one side of a given interaction interface, and dozens to hundreds of designs must be screened to uncover and validate binders acting by the desired mechanism^32,33^.

Biomolecular condensate formation is fundamentally driven by networks of multivalent interactions^11,28,34–37^ (**Fig. 1A**). In proteins, these multivalent interactions frequently involve globular domains^28,38–40^ as well as IDRs. Therefore, in principle, ligands that bind interaction interfaces of globular domains could be employed to both study and control condensate formation. Monovalent ligands could act as inhibitory ‘chain terminators’ of the multivalent interaction network, and bridging ligands could conversely enhance phase separation by strengthening multivalent interactions through non-covalent cross-links^35^.

**Figure 1.**
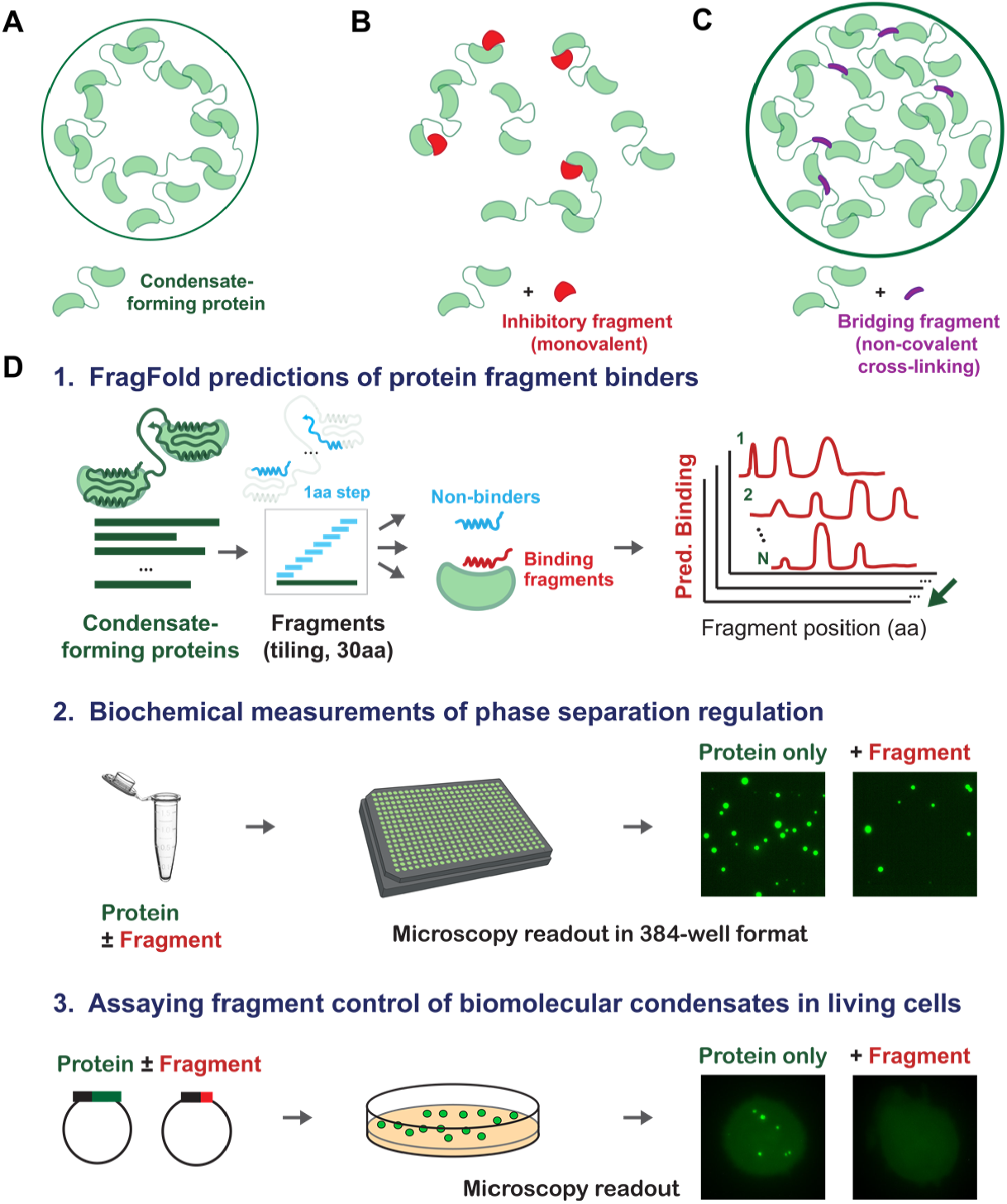
Protein fragments as regulators of phase separation. (**A-C**) Conceptualization of protein fragments as general regulators of protein condensates, following the nature of condensates as networks of multivalent interactions (**A**). (**B**) Monovalent binder fragments of condensate-forming proteins should be able to act as inhibitors of condensates by terminating networks of multivalent interactions. (**C**) Bridging fragments that form interactions across a multivalent interface without blocking it should be able to act as bridging ligands that stabilize and enhance condensate formation. (**D**) Schematic of the approach introduced and employed in this work. 1. FragFold^41^ is used to predict fragments of condensate-forming proteins that can form interactions with their full-length parent protein, which are predicted to act as condensate modulators. 2. – 3. *In vitro* and in-cell microscopy assays of condensate formation allow assessment of the function of FragFold-designed condensate modulators.

We reasoned that protein fragments^41,42^ derived from full-length proteins should provide a practical realization of the theoretical concepts of monovalent and bridging ligands. Protein fragments could hence provide a broadly applicable approach to study, manipulate, and therapeutically target diverse protein condensates. Further, such fragments would directly employ native interactions, shedding light on drivers of condensate formation. Fragments that block multivalency-promoting interfaces would then be expected to act as generalizable condensate inhibitors (**Fig. 1B**). Conversely, fragments that promote multivalent interactions by bridging proteins should act as condensate enhancers (**Fig. 1C**). Consistent with this idea, a retro-inverso peptide derived from a dimerization domain of SARS-CoV-2 nucleocapsid was recently found to inhibit nucleocapsid condensate formation^43^. Recently, it has become possible to discover protein fragments to regulate protein interactions on a massively parallel scale using AI methods^41^. Complementary AI-driven predictions and high-throughput experiments have demonstrated that protein fragments are generalizable inhibitors of diverse proteins^41,42,44–46^.

Here, we show that protein fragments which inhibit or enhance phase separation can be readily discovered by an AI-driven algorithm, FragFold^41^, that predicts binders of target proteins. We demonstrate that these fragments are functional both *in vitro* and in living mammalian cells (**Fig. 1D**), achieving an overall success rate of 50% across diverse phase-separating proteins. Our findings have major implications for the study of fundamental biology of biomolecular condensates, as well as the therapeutic targeting of condensates associated with disease.

## Results

### Protein fragments are generalizable regulators of biomolecular condensates

We sought to establish the functionality and generalizability of protein fragments as regulators of biomolecular condensates of diverse proteins. To achieve this goal, we employed FragFold – a massively parallel AI method for the discovery of protein fragment binders^41^ shown to predict inhibitory fragments with high accuracy and precision^41,42^. For each phase-separating protein, we applied FragFold to predict binding between all possible protein fragments and the full-length parental protein, reasoning that fragment binders have a high chance of either inhibiting or enhancing multivalent interactions (**Fig. 1A**). Indeed, recent work supports the idea that most interface-binding protein fragments predicted by FragFold regulate biologically relevant binding interactions^41,42^. Notably, we employed no knowledge of specific interfaces that were known or suspected to be involved in multivalent interactions driving phase separation in the predictions or their evaluation. Instead, we selected a set of fragments corresponding to the strongest and broadest local maxima in FragFold-predicted binding for each protein – following our previous systematic evaluation of the correlation between these predictions and massively parallel measurements of protein fragment inhibitory activity^41^. Overall, we computationally screened 2,235 protein fragments and selected 18 (3-5 per protein) for experimental investigation.

We first employed FragFold to design protein fragment regulators of three phase-separating proteins with diverse structures, cellular localizations, and biological functions (**Fig. 2**): Ras GTPase-activating protein-binding protein 1 (G3BP1), the core protein mediating the formation of stress granules^47,48^ (**Fig. 2A-B**); the SARS-CoV-2 nucleocapsid protein (SARS2-NCAP or NCAP), which forms condensates with an important role in the viral replication cycle^49^ (**Fig. 2C-D**); and TAR DNA-binding protein (TDP-43), which modulates splicing, transcription, and translation^50^, and forms both phase-separated droplets and pathological aggregates in ALS and Alzheimer’s disease^51,52^ (**Fig. 2E-F**). All three of these proteins and their biomolecular condensates are of considerable therapeutic interest. Each of these proteins binds RNA; G3BP1 and TDP-43 condensate formation is enhanced by RNA, whereas SARS2-NCAP phase separation requires RNA. In comprehensive FragFold predictions of fragment binding, each protein yielded numerous fragments with strong predicted binding activity. For each protein, we selected a set of fragments (five per protein) corresponding to FragFold-predicted binding peaks (**Fig. 2A,C,E**, *Methods*).

**Figure 2.**
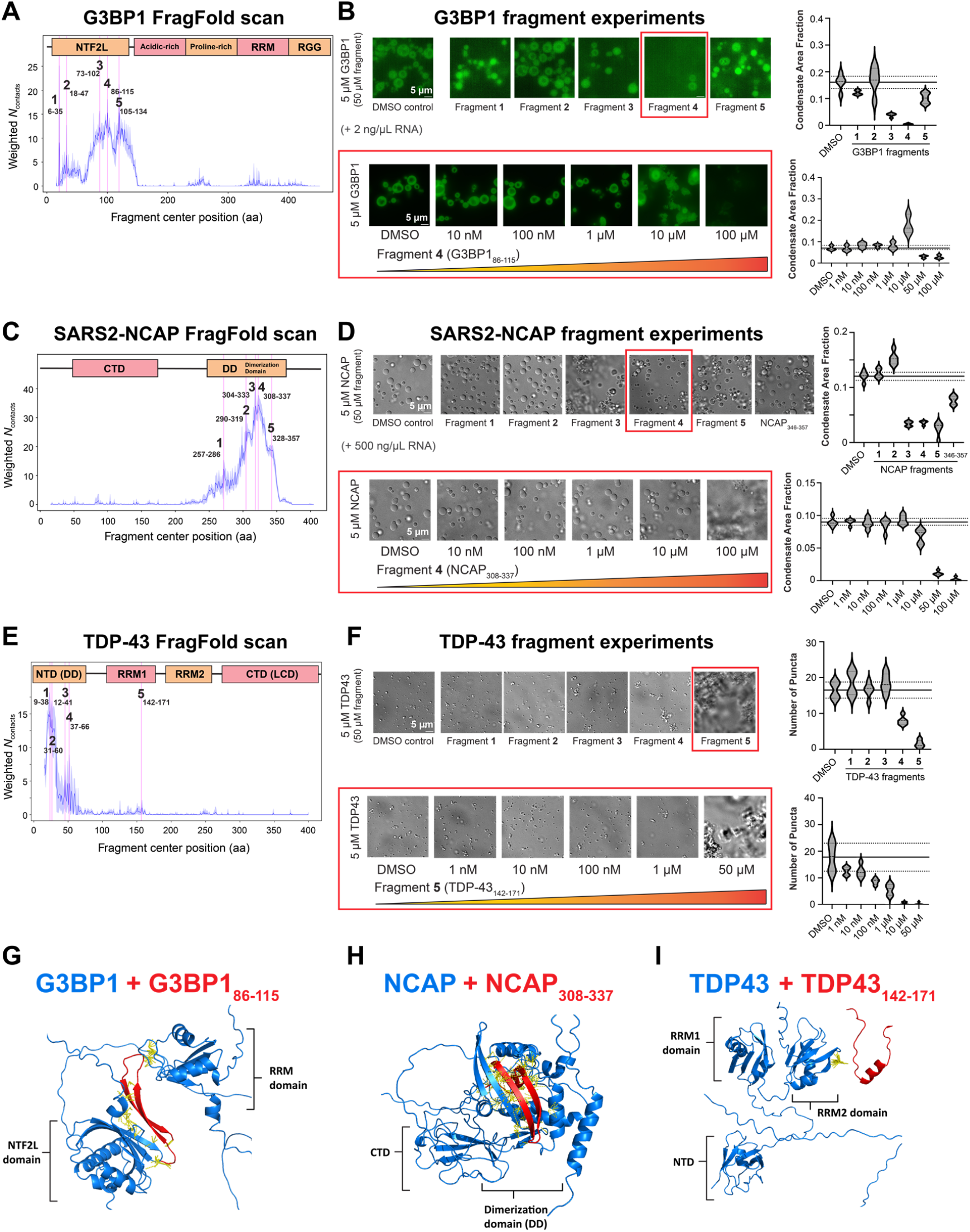
FragFold discovery of protein fragment regulators of condensate formation generalizes across diverse proteins. (**A**) – (**F**): FragFold-predicted protein fragment binding profiles to full-length phase-separating proteins (30 aa fragments, 1 aa step size), *left*, and corresponding experimental effects of chemically-synthesized protein fragments on biomolecular condensates of the parental protein *in vitro, right*, for: (**A**) – (**B**) G3BP1-GFP, in the presence of 2 ng/µL RNA; (**C**) – (**D**): SARS-Cov-2 nucleocapsid (SARS2-NCAP, or NCAP), in the presence of 500 ng/µL RNA; (**E**) – (**F**): TDP-43. In each case, the fragments selected for synthesis and experimental testing are indicated overlaid on the fragment scan plots ((**A**), (**C**), (**E**)), and a single fragment with a condensate-inhibitory effect (red square outline) is highlighted with its corresponding FragFold binding model ((**G**), (**H**), (**I**)). (**B**), (**D**), (**F**), *bottom*: Titrations of selected phase-separation-modulating protein fragments, as indicated. (**B**), (**D**), (**F**), *right*: quantification of measurements for each fragment and titrations of example fragments. Mean ± S.D. of the DMSO control is indicated with horizontal solid (mean) and dashed (± S.D.) lines across each plot. DMSO, DMSO control.

We chemically synthesized the FragFold-predicted protein fragment binders and tested their effects on condensate formation *in vitro* (**Fig. 1D, Fig. 2B,D,F**, *Methods*). Since protein fragments are most likely to bind using native-like interactions^41,42^, presumably with similar interaction strengths, we initially tested a 10-fold molar excess of fragments. For each protein, our approach readily generated fragments that strongly inhibit condensate formation (**Fig. 2B,D,F**, *upper*; **Fig. S1**), with no modification required from the genetically encodable fragment sequences. One example fragment of each protein was further investigated in titration experiments (**Fig. 2B,D,F**, *lower*).

In the case of G3BP1 mixed with 2 ng/µL RNA, two of five fragments tested substantially inhibited condensate formation (**Fig. 2B**, fragments 73-102 and 86-115; here and throughout, referring to the amino acid residues of the full-length protein spanned by the fragment). These reductions were statistically significant (*p* < 0.01, Student’s t-test). G3BP1_86-115_, the fragment with the strongest inhibitory effects, is both derived from and predicted to bind to the NTF2L dimerization domain. FragFold predicts the fragment binds G3BP1 in a native-like manner mimicking NTF2L dimerization^53^ (**Fig. 2G, Fig. S2A**). Our experimental results and FragFold structural model are thus congruent with this fragment acting as a monovalent ligand inhibitor (**Fig. 1B**). The titration of G3BP1_86-115_ also captured an intermediate state at 10 µM fragment : 5 µM protein where condensates appeared to be enlarged but more diffuse, likely representing a partially destabilized state (**Fig. 2B**). NTF2L dimerization is essential for G3BP1 phase separation *in vitro* and stress granule formation in cells^54^, and the NTF2L domain is targeted by nsP3 viral protein that disrupts stress granule formation^55^. Our systematic, AI-driven approach successfully identified NTF2L domain dimerization as essential for condensate formation.

In the case of SARS2-NCAP, condensate formation requires both NCAP and RNA^56^. Here we included 500 ng/µL RNA. We found that three of five fragments discovered with FragFold – NCAP_304-333_, NCAP_308-337_, and NCAP_328-357_ – strongly and significantly inhibited condensate formation (**Fig. 2D;** *p* < 10^−4^, Student’s t-test). NCAP_308-337_ inhibits condensate formation even at a 2:1 ratio (**Fig. 2D**, *lower*). FragFold models of fragment binding mimic the experimentally determined swapped-dimer structure of the dimerization domain (DD) of SARS2-NCAP^57^ (**Fig. 2H** and **Fig. S2B**). Our experimental results and FragFold structural models are therefore consistent with NCAP_308-337_ acting as a monovalent ligand inhibitor (**Fig. 1B**). Notably, these results demonstrate that protein fragments can inhibit condensates that require both protein and RNA for their formation.

Previously, the SARS2-NCAP DD was targeted using rational design of retro-inverso peptides^43^, yielding a peptide covering residues 346-357 that inhibits condensate formation. All other rationally designed peptides based on the DD were not functional in this prior study. Here, we found that FragFold AI-designed protein fragments – with no rational or structure-guided steps in design or testing – converged on the DD interface, and 3 of 5 fragments tested inhibited phase separation (**Fig. 2D**). In addition, fragment 290-319 from the DD appears to weakly enhance condensate formation (**Fig. 2D**), increasing the number of condensates at the expense of condensate size (**Fig. S1B**). The L-amino acid peptide NCAP_346-357_ mirroring the retro-inverso peptide^43^ was also inhibitory (*p* < 10^−3^, Student’s t-test), consistent with previous measurements, but measured inhibition was weaker than that of the FragFold designs. The fragment with the strongest contacts in FragFold models, NCAP_308-337_, was the strongest inhibitor (**Fig. 2D**). Thus, in the case of SARS2-NCAP, our AI-driven approach outperforms rational design in identifying interaction-inhibitory fragments.

In the case of TDP-43, we found that two of five fragments discovered with FragFold – TDP-43_37-66_ and TDP-43_142-171_ – strongly and significantly inhibited condensate formation (**Fig. 2E-F;** *p* < 0.05, *p* < 0.01, Student’s t-test). TDP-43_37-66_ maps to the N-terminal globular oligomerization domain which is known to be required for condensate formation^58^. FragFold predicts that TDP-43_37-66_ binds full-length TDP-43 in a native-like dimerization interaction (**Fig. S2C**), consistent with this fragment blocking this key oligomerization interface driving phase separation. TDP-43_142-171_ was the stronger inhibitor, beginning to inhibit condensate formation by a 1:50 ratio of fragment : protein. The FragFold model for TDP-43_142-171_ binding indicates that it binds the RNA-binding motif RRM2 (**Fig. 2I, Fig. S2D**), with TDP-43_142-171_ itself sourced from RRM1. Consistent with our results, RRM domain interactions play a major role in TDP-43 phase separation, including in the absence of nucleic acids^59–61^, the conditions employed in our experiments.

### AI-discovered protein fragments readily inhibit or enhance phase separation of focal adhesion kinase

We next applied FragFold to design protein fragments regulating the phase separation of focal adhesion kinase (FAK), a mammalian protein which promotes cell growth and motility^62,63^ and is upregulated in multiple cancers^64,65^. FAK is therefore a target of considerable interest in cancer therapeutics^66,67^, and the role of FAK phase separation in physiology and disease is an active area of inquiry^68–70^. Compared to G3BP1, SARS2-NCAP, and TDP-43, the protein interactions driving FAK condensate formation are less understood, providing an opportunity to discover these interactions using protein fragments.

We used FragFold to computationally screen all possible 30-amino acid fragments of FAK, selected three fragments for chemical synthesis (**Fig. 3A**), and experimentally assessed their effects on GFP-FAK condensate formation (**Fig. 3B**). Two of the three fragments significantly impacted phase separation. FAK_379-408_ strongly inhibited condensate formation, while FAK_977-1006_ enhanced condensate formation (**Fig. 3B-C**). By contrast, an unrelated fragment of the bacterial protein FtsZ^42^ (FtsZ_34-63_) had no effect on phase separation (**Fig. S1C**). Unlike the condensate-regulatory protein fragments we discovered for G3BP1, SARS2-NCAP, and TDP-43, FAK_977-1006_ strongly enhanced condensate formation and resulted in significantly larger droplets (**Fig. 3B, Fig. S1C**). This behavior is consistent with a bridging fragment increasing multivalent interactions (**Fig. 1C**). FAK_977-1006_ is predicted to bind to a helical bundle formed by the FAT domain, with the fragment sourced from this same domain (**Fig. 3D**, *right*).

**Figure 3.**
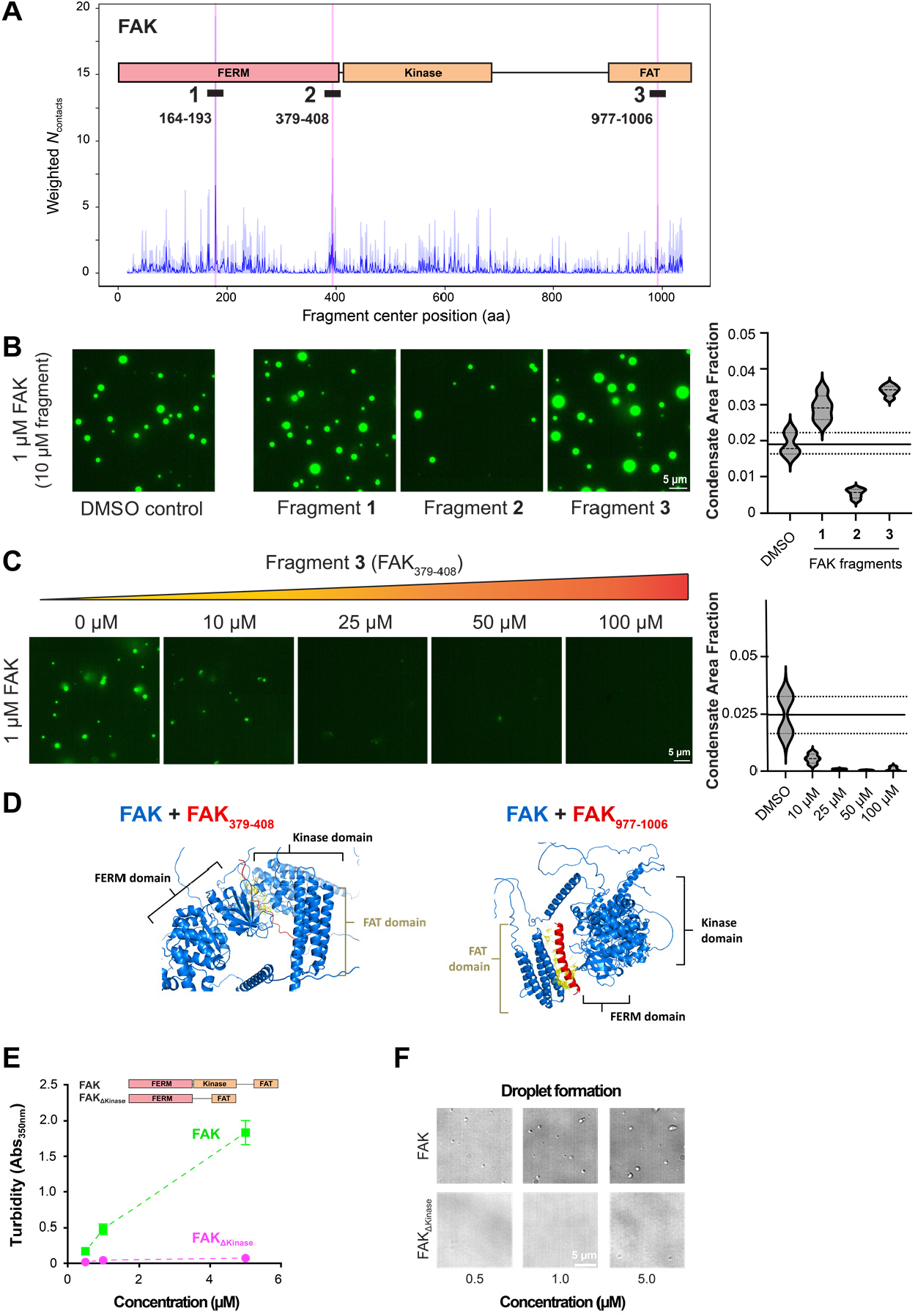
FAK protein fragments robustly regulate FAK condensates and reveal role of the kinase domain in condensate formation. (**A**) FragFold prediction of FAK protein fragments that bind to FAK. The domain structure of FAK and the fragments selected for synthesis and experimental testing are highlighted. (**B**) *Left*: *In vitro* microscopy results for condensate inhibition by FAK fragments. In all cases, FAK is present at 1 µM and fragment (if present) is added at 10 µM. *Right*: Quantification of the results to the left. Mean ± S.D. of the DMSO control is indicated with horizontal solid (mean) and dashed (± S.D.) lines across each plot. (**C**) *Left*: Titration of FAK condensates with increasing concentrations of FAK_379-408_. The visual difference in the puncta compared to (B) is due to the higher total DMSO concentration matching the DMSO in up to 100 uM fragment added. *Right*: quantifications of the results to the left, as in (B). (**D**) FragFold models for the binding modes of fragments FAK_379-408_ and FAK_977-1006_. (**E**) Turbidity measurements and (**F**) droplet formation assay for condensate formation by full-length FAK and FAK with the kinase domain deleted (FAK_ΔKinase_).

These structural predictions are consistent with two models for how FAK_977-1006_ could enhance condensate formation. First, the FAT domain is involved in dimerization interactions of FAK^71^, which could contribute to condensate formation. FAK_977-1006_ binding could enhance dimerization by modulating FAT dimer interactions without blocking their binding interface, consistent with the FragFold model overlaid with an experimental structure of the dimer^72^ (**Fig. S2E**). Second, the FAT domain can also interact with the FERM domain of FAK^73^, providing multivalency that contributes to condensate formation. The FAT-FERM interaction is enhanced by paxillin LD4^73^, which binds the FAT domain in a similar mode to FAK_977-1006_ (**Fig. S2F**). Indeed, the FragFold structural model shows the fragment bridging the FAT and FERM domains.

Titrating the inhibitory fragment FAK_379-408_, we observed a near-total abrogation of condensate formation by a fragment:protein ratio of 25:1 (**Fig. 3C**). Thus, FAK_379-408_ acts as a potent inhibitor of condensate formation. However, FAK_379-408_ did not dissolve pre-formed FAK condensates (**Fig. S2G**), suggesting inhibitory fragments are not sufficient to disrupt mature FAK condensates under the specific *in vitro* conditions tested.

The FragFold-predicted binding mode of FAK_379-408_ provided new insight into the interactions driving phase separation (**Fig. 3D**, *left*). We found that FAK_379-408_ is predicted to bind at an interface between the FERM and kinase domains of FAK. In practice, the fragment likely binds to either the FERM or the kinase domain at any one time with FragFold overpredicting simultaneous interactions, as seen previously^41^. The FERM-kinase domain interaction is thought to regulate FAK function intramolecularly^74^ (and sequesters regulatory phosphosite Tyr397), and our results suggested this interaction also contributes to the intermolecular interaction network in FAK condensates.

### The kinase domain of FAK is required for condensate formation, consistent with FragFold inhibitory fragment mapping

To experimentally test if the FERM-kinase domain interaction is important for condensate formation, we purified recombinant FAK with the kinase domain deleted. We previously demonstrated that FAK kinase activity is not required for phase separation, since ATP is not required for condensate formation and unphosphorylated FAK phase separates similarly to phospho-FAK^71^. We first measured phase separation by measuring solution turbidity at 350 nm^68^. While full-length FAK undergoes a concentration-dependent increase in turbidity, as previously reported^68^, the kinase deletion mutant exhibited no increase in turbidity even at 5 µM (**Fig. 3E**). We confirmed the mutant was not forming visible condensates with DIC microscopy (**Fig. 3F)**. These experimental results are consistent with the FragFold structural model (**Fig. 3D**) and fragment assays (**Fig. 3B-C**) and suggest that the kinase domain participates in multivalent interactions promoting phase separation. Thus, inhibitory fragment mapping with FragFold successfully revealed that the kinase domain is required for condensate formation.

### FAK fragments control phase separation in living mammalian cells

Since FAK_379-408_ acts as a strong inhibitor of condensate formation (**Fig. 1B**) *in vitro*, we next tested the ability of FAK_379-408_ to regulate FAK condensates in a cellular context. We used a previously validated assay to induce FAK cytosolic condensates by overexpressing FAK^71^. Mouse embryonic fibroblasts (MEFs) were co-transfected with plasmids encoding GFP-tagged FAK and FAK_379-408_ fused to DsRed by a co-translationally cleaved 2A peptide^75,76^. We imaged cells by epifluorescence microscopy and quantified the number of cytosolic condensates per cell (*Methods*, **Fig. 4A**). Cells overexpressing GFP-FAK formed cytosolic condensates as expected^71^ (**Fig. 4B**). However, cells expressing both GFP-FAK and FAK_379-408_ formed substantially fewer condensates (**Fig. 4B-D**), and condensate formation depended on the relative stoichiometry of GFP-FAK and FAK_379-408_ expressed in each cell. Quantifying the number of condensates per cell across 128 co-transfected cells, we found that an increase in the fragment:FAK intensity ratio from ≤ 0.5 to ≥ 1 dramatically decreases the number of condensates per cell, from 8.2 ± 3.0 (mean ± standard error) to 0.6 ± 0.2 – with a significant negative correlation between the number of condensates and the intensity ratio (Spearman *ρ* = −0.54, *P* < 10^−10^; **Fig. 4D-E**). Together, these results demonstrate that FAK_379-408_ suppresses FAK condensate formation in living mammalian cells in a dose-dependent manner.

**Figure 4.**
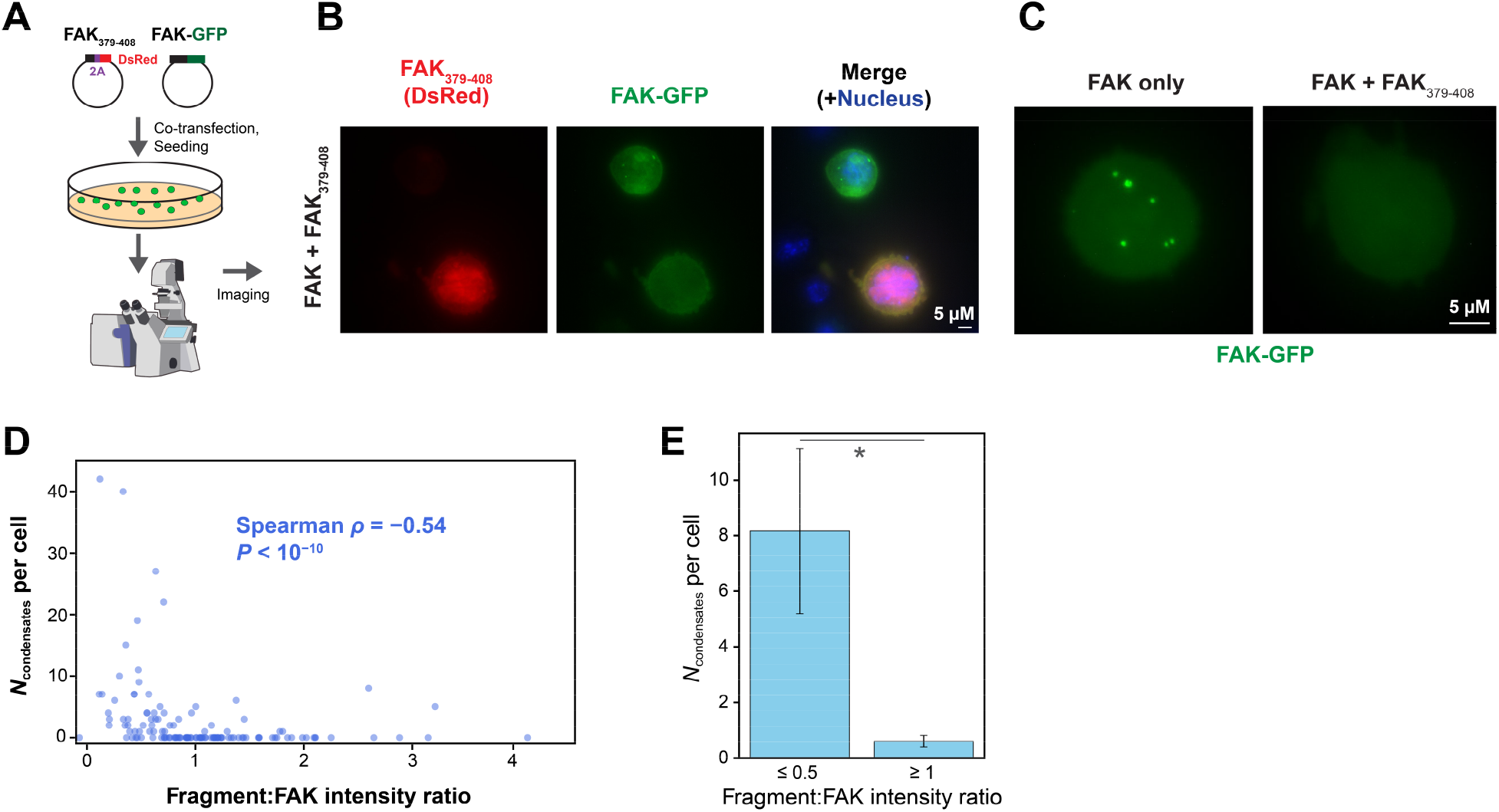
FAK fragment 379-408 inhibits FAK condensates in living cells. (**A**) Mouse embryonic fibroblasts (MEFs) were co-transfected with plasmids for expression of FAK_379-408_-2A-DsRed and FAK-GFP. After 24 hours the cells were seeded onto poly-D-Lysine plates and imaged for DNA (Hoechst), DsRed, and GFP fluorescence. (**B**) DNA, FAK_379-408_, and FAK imaging. Cells expressing significant FAK-GFP but not FAK_379-408_ form condensates; cells expressing both FAK-GFP and FAK_379-408_ exhibit drastic reduction of FAK condensates. (**C**) Representative cell expressing FAK-GFP only vs. cell expressing FAK-GFP + FAK_379-408_-2A-DsRed, at the same FAK transfection level (GFP intensity of ∼250 a.u.; *Methods*). (**D**) Quantification of microscopy results. Number of FAK condensate puncta (*N*_condensates_) per cell as a function of the fragment:FAK (DsRed:GFP) intensity ratio in co-transfected cells. The number of puncta is negatively correlated with the intensity ratio (Spearman *ρ* = −0.54, *P* < 10^−10^). (**E**) Bar plot from data in (D), comparing number of condensates per cell for fragment:FAK intensity ratios of ≤ 0.5 and ≥ 1. Bar height, mean value; error bars, standard deviation. *, difference is significant at *p* < 0.05, Student’s t-test.

## Discussion

In this work, we have demonstrated that protein fragments can act as generalizable and robust regulators of biomolecular condensates, and that such fragments can be readily discovered and designed using AI methods. Our approach allows for the design of protein fragments that inhibit or enhance the phase separation of a specific target protein. Each fragment was designed to bind its target protein, and the inhibitory fragments did not share common physicochemical features (**Table S1**). Until now, the field has lacked generalizable methods to inhibit the formation of specific condensates. Thus, protein fragments are a powerful experimental tool that can be used test necessity and sufficiency of phase separation for biological functions. Protein fragments and FragFold-predicted binding modes can also be used to experimentally determine specific interactions driving phase separation, and this approach enabled the identification of a previously unknown interdomain interaction required for FAK condensate formation.

Our approach is effective across diverse phase-separating proteins, and the corresponding fragments are functional both *in vitro* and in living cells, with no modifications required from the linear, genetically encodable peptide sequence. The proteins investigated in this work, and their condensates, are also of considerable biomedical interest, including in the context of cancer^64,65^ and neurodegeneration^51,52^. The regulatory fragments reported here thus provide a promising foundation for developing new therapeutics targeting biomolecular condensates.

To date, FragFold has been demonstrated to robustly predict purely globular protein interactions and globular-IDR interactions (e.g., FtsZ C-terminal IDR binding)^41^; consistent with these results, FragFold-designed condensate modulators reported here all take advantage of interactions where at least one partner is globular. However, even for proteins with roles for IDR interactions in their phase separation (e.g., G3BP1^77^, TDP-43^78^), we find fragment-driven modulation of globular interactions can suffice to control condensate formation.

Unmodified protein fragments might be expected to function at similar concentrations to their targets, due to inherently similar native-like interactions. Since we used micromolar protein concentrations to form condensates *in vitro*, this typically yields micromolar-range inhibitory concentrations. Looking forward, for therapeutic applications fragments would often benefit from further engineering, such as affinity maturation through deep mutational scanning^41^. A related consideration is aggregation. Indeed, we found several fragments investigated here aggregate at high concentrations (**Fig. S3**). However, very little aggregation is observed for FAK_379-408_, and both FAK_379-408_ and TDP-43_142-171_ are active at concentrations well below those triggering aggregation (**Fig. 2, Fig. 3**). Further, for G3BP1_86-115_, fragment aggregates do not colocalize with the protein target, so aggregation is apparently distinct from inhibition (**Fig. S3E**). In future work aggregation may be circumvented by engineering higher affinity^41^, developing fragment-inspired *de novo* binders^33^, or computationally removing aggregation-driving features^33,79^.

AI-driven discovery of protein fragments is deeply synergistic with recently developed high-throughput experimental methods^23,24^. Our results immediately suggest that such assays can be adapted to massively parallel screening for condensate-inhibitory and -enhancing functionalities of large libraries of protein fragments^24^. The exciting possibilities for future work in this direction are underscored by the high yield of functional protein fragments reported here.

More broadly, our results extend the ability of FragFold to discover protein fragments as universal regulators of protein interactions across species from bacteria^41^ to human. Protein fragments regulate diverse protein interactions, including those driving cell growth and division^41^ and condensate formation. These results underscore that protein fragments are broadly functional biological entities – with immense promise as bioengineering tools to study and control protein interactions, and cellular physiology, across species and systems.

## Acknowledgements

We thank members of the Li and Case labs for helpful discussions. This work was supported by awards K99 GM148718 (to A.S.), R35 GM124732 (to. G.W.L.), DP2 GM149549 (to L.B.C.), a Ludwig Center Postdoctoral Fellowship from MIT’s Koch Institute for Integrative Cancer Research (to J.S.), a National Cancer Center Postdoctoral Fellowship (to J.S.), and an NSF GRFP fellowship (to J.D.R.). K.J.W. and J.D.R. were supported by training grant T32 GM136540. G.W.L. is an Investigator of the Howard Hughes Medical Institute.

## Author Contributions

Initial research conceptualization was performed by A.S. and L.B.C. Further conceptualization was performed by A.S., J.S., and L.B.C. Research design was performed by A.S. and J.S. Computational discovery of condensate-regulatory protein fragments was performed by A.S. Experiments were performed by J.S. with feedback from A.S. and L.B.C. Protein purifications were performed by J.S., K.J.W., and J.D.R. Data analysis was performed by A.S. and J.S. The manuscript was written by A.S. and J.S., with feedback from G.W.L. and L.B.C. and approval of all authors.

## Resource Availability

### Lead Contact

Requests for resources and reagents should be directed to the lead contacts, Andrew Savinov and Lindsay Case.

### Materials Availability

The reagents and materials used in this study are available upon reasonable request.

### Data and code availability

The code and software used in this paper – and to run FragFold protein fragment predictions for any protein of interest – is available on GitHub (https://github.com/gwlilabmit/FragFold). Data are provided in the main text, the Supplementary Materials, and a Source Data file available via figshare (https://doi.org/10.6084/m9.figshare.32224497).

## Supplementary Materials

### Methods

#### FragFold design of biomolecular-condensate modulating protein fragments

To design protein fragments to control protein biomolecular condensate formation, FragFold^41^ was used to predict binding of every possible 30-amino acid protein fragment of each condensate-forming protein of interest to its parental full-length protein. FragFold predictions were run using the published FragFold software (https://github.com/gwlilabmit/FragFold) as previously described^41^. Inputs for the FragFold predictions were the full-length parental protein sequences from the corresponding UniProt entries, as follows: FAK, P34152; SARS2-NCAP, P0DTC9; G3BP1, Q13283; and TDP-43, Q13148. These same sequences were used by FragFold to automatedly generate the fragments and predict their binding interactions. Other inputs were the default running parameters for the FragFold software (e.g., fragment_length: 30, fragment_ntermres_start: 1, protein_copies: 1, protein_ nterm_res: 1). Fragments were selected for synthesis and experimental testing based on selection of local maxima (peaks) in the FragFold fragment tiling profile of weighted *N*_contacts_ vs. fragment sequence position (e.g., **Fig. 2A**), starting from global maxima for each phase-separating protein investigated; and also prioritizing peaks with a broader peak width (consisting of a larger number of contiguous binding fragments), as previously demonstrated to give good agreement with massively parallel measurements of protein fragment inhibitory function in living cells^41,42^. Selected fragments were then synthesized and experimentally tested as described in the following sections.

#### Protein fragment sequences and synthesis

Protein fragments of FAK, SARS2-NCAP, G3BP1, and TDP-43 were synthesized at the Biopolymers & Proteomics Laboratory at the MIT David H. Koch Institute for Integrative Cancer Research. Synthesis was performed using an Intavis Model MultiPep synthesizer at a 5 mg scale and 80-85% purity. We confirmed that the results for an example fragment, FAK_379-408_, were unchanged when the fragment was further HPLC-purified to 95% purity (**Fig. S4**). Fragment sequences were derived from UniProt entries for each protein as follows: FAK, P34152; SARS2-NCAP, P0DTC9; G3BP1, Q13283; and TDP-43, Q13148. The FtsZ control protein fragment (FtsZ_34-63_Savinov2022) was synthesized as described previously^41^. Fragment sequences are listed in **Table S1**.

#### 384-well in vitro microscopy assay for condensate formation and modulation by protein fragments

384-well glass-bottomed plates (Brooks) were incubated in preheated 5% Hellmanex III (∼65°C) for 3–4 h with periodic inversion to remove trapped air bubbles and thoroughly rinsed with MilliQ H_2_O. Plates were then washed with 1 M NaOH for 1 h at room temperature, washed again with MilliQ H_2_O and dried thoroughly in a fume hood. Wells were incubated with 50 µL of 20 mg/mL mPEG-silane in 95% ethanol overnight at room temperature in the dark. The following day, the plate was thoroughly rinsed with MilliQ H_2_O, dried, sealed with foil, and stored until use. On the day of imaging, required wells were opened and passivated with imaging buffer containing 1 mg/mL BSA for at least 30 min. After buffer removal, 50 µL protein mixtures were added, and samples were imaged by fluorescence microscopy. Protein stocks were diluted in their respective storage buffer before being diluted into an experimental buffer that yielded the desired final concentrations of all buffer components (buffer, salt, etc.) after accounting for contributions from the protein storage buffers. Buffer conditions used for each protein were as follows:

FAK: 1 μM FAK, 50 mM HEPES (pH 7.4), 50 mM KCl, 1 mg/mL BSA, 1 mM TCEP. Fragments were added to a final concentration of 10 μM (10:1 fragment:protein ratio) unless noted otherwise.

SARS2-NCAP: 5 μM SARS2-NCAP, 50 mM Tris-HCl (pH 7), 150 mM NaCl, 5 mM MgCl_2_, 1 mM DTT, 1 mg/mL BSA, 5% PEG, 0.5 ug/µL Poly (A) RNA (Cytiva 27-4110-01). Fragments were added to a final concentration of 50 μM (10:1 fragment:protein ratio) unless noted otherwise.

G3BP1: 5 µM G3BP1, 50 mM HEPES (pH 7.4), 50 mM KCl, 5 mM MgCl2, 1 mM TCEP, 1 mg/mL BSA, 2 ng/µL FLuc RNA. Fragments were added to a final concentration of 50 μM (10:1 fragment:protein ratio) unless noted otherwise.

TDP-43: 5 μM TDP-43, 20 mM HEPES (pH 7.4), 150 mM NaCl, 1 mM DTT, 0.3 units/µL TEV protease (NEB, P8112S). Fragments were added to a final concentration of 50 μM (10:1 fragment:protein ratio) unless noted otherwise.

For FAK and TDP-43, buffer was added first, and proteins required for phase separation were added last. For G3BP1 and SARS2-NCAP, proteins were incubated with peptides for 15 minutes before adding the RNA to induce phase separation. Samples were mixed by pipetting and loaded into 384-well PEG-silane–coated glass-bottom plates. Samples were incubated for 30-60 min before imaging. For each experiment, regions of interest were randomly selected for imaging, and all images were acquired within 10 minutes of each other.

#### Cell culture, plasmids and transfection

Mouse embryonic fibroblast (MEF ATCC CRL-2645) cells were maintained in Dulbecco’s Modified Eagle Medium (DMEM; high glucose, Invitrogen) supplemented with 10% fetal bovine serum (FBS, Thermo Fisher Scientific), 1% penicillin–streptomycin, and 2 mM Glutamax at 37°C in a humidified incubator with 5% CO_2_. MEFs were passaged using TrypLE Express (Thermo Fisher Scientific) according to manufacturer’s protocol.

For in-cell experiments, FAK_379-408_ was C-terminally fused in-frame with a 2A peptide sequence from porcine teschovirus, followed by DsRed, allowing co-translational cleavage of the protein fragment from the DsRed protein by the “StopGo” mechanism as they are synthesized^75,76^ (Twist Biosciences). The gene fragment was cloned into a mammalian expression vector pMEGFP-C1 using NheI and MfeI (New England Biolabs). The construct was verified by long read nanopore sequencing (Plasmidsaurus).

Transfections with GFP-FAK and FAK_379-408_-2A-DsRed were performed using Lipofectamine− 3000 (Invitrogen) with minor modifications to the manufacturer’s protocol. Briefly, cells were seeded in 6-well plates to reach ∼50–70% confluency at the time of transfection. For each well, 5 µg of plasmid DNA was diluted in 125 µL Opti-MEM containing 10 µL P3000 reagent and incubated for 5 min at room temperature. In parallel, 7.5 µL Lipofectamine 3000 was diluted in 125 µL Opti-MEM. The Lipofectamine mixture was then combined with the DNA mixture and incubated for an additional 5 min at room temperature. The resulting complexes were added dropwise to cells in complete growth medium, followed by gentle swirling to ensure even distribution. Cells were incubated at 37°C overnight. After 24 hours, cells were trypsinized and 50,000 cells were seeded into 8-well chambers (iBidi) coated with poly-D-Lysine (Thermo Fisher Scientific). 15 minutes after seeding, cells were imaged by epifluorescence and differential interference contrast (DIC) microscopy at 37°C with 5% CO2.

#### Microscopy

Epifluorescence and differential interference contrast (DIC) images were acquired using LAS-X acquisition software controlling a Leica DMi8 microscope equipped with DIC optics and a Plan Apo 100×/1.47 NA TIRF objective. Illumination was provided by an LED8 light source with two interchangeable filter cubes (excitation at 391/32, 479/33, 554/24, 638/31 or 473/22, 539/24, 641/78, 810/80). Images were collected using a Hamamatsu Flash 4.0 V3 CMOS camera.

Representative images of *in vitro* condensates were captured using epifluorescence or DIC microscopy. Buffer conditions for individual experiments are indicated in the figure legends. Fluorescence images were analyzed using Fiji (ImageJ). All images were acquired using identical microscope and camera settings. Images were converted to 8-bit format. Images were thresholded using the IsoData thresholding algorithm with a dark background, applying a fixed intensity threshold range of 30–80 uniformly across all images. Thresholded images were converted to binary masks, and particles were identified using the Analyze Particles function with a minimum particle size cutoff of 0.05 pixel units and no upper size limit.

Particles of G3BP1 and NCAP condensates were analyzed using CellPose^80,81^ (v4.0.8) in the graphical user interface on macOS (python 3.10). Segmentation was performed using pretrained *cyto* model (CPSAM, cellpose V4). Images were processed as 2D inputs using automatic intensity normalization with default parameters, unless otherwise specified. Segmentation masks were exported as PNG images. PNG masks were analyzed in Fiji (ImageJ). Images were converted to 8-bit format and thresholded using the Huang method. For G3BP1 condensates, particles were identified using the Analyze Particles function with a minimum particle size cutoff of 0.5 pixel units. For NCAP condensates, particles were identified with a minimum size threshold of 2000 pixel units. Particles of TDP-43 were too small for quantification by CellPose; therefore, random fields of view were selected for manual particle counting.

#### Turbidity assays of condensate formation

Unlabeled proteins were diluted in 25 mM Hepes (pH 7.0), 50 mM KCl, 1 mM TCEP, 0.1% (1 mg/mL) BSA. After a 45–75 min incubation, solution was transferred to Quartz cuvette and Absorbance at 350 nm was measured in a Spectrophotometer (Agilent or Thermo Fisher Scientific). While we use 1 mg/mL BSA (0.1%) to prevent nonspecific interactions with surfaces, we note that this is well below the BSA concentration necessary to induce crowding (typically 100 mg/mL or greater).

#### In-cell condensate puncta and fluorescence intensity analysis

Cells transfected with GFP-FAK and FAK_379-408_-2A-DsRed were plated on poly-D-Lysine as described. Tile scans covering 15 × 15 field of view (FOV, 132 × 132 micron each) were imaged with epifluorescence microscopy, with Z-stacks of each FOV to capture the entire volume of cells. Maximum intensity projections of each FOV were created, and the FOVs were stitched using LASX software. Images were analyzed using a custom macro as follows: For each cell, a fixed 600 × 600 pixel region of interest (ROI) was manually selected and added to the ROI manager. The same set of ROIs were applied to all channels to make sure that measurements were obtained from same cells across channels. For each ROI, macro generated a new image stack in which each slice corresponds to one cell. Mean fluorescence intensity of each cell was measured using ImageJ “Analyze Particles”. The number of puncta in each cell was counted manually.

#### Protein expression and purification

##### Focal adhesion kinase

Full length Focal adhesion kinase (FAK) protein was purified as described previously^71^. Briefly, His-mEGFP-FAK was expressed using a baculovirus system in HighFive (Thermo) *Trichoplusia ni* cells. Cells were harvested by centrifugation and lysed using a dounce homogenizer on ice in lysis buffer containing 25 mM HEPES (pH 7.5), 30 mM imidazole (pH 7.5), 500 mM NaCl, 10% glycerol, 5 mM βME, and cOmplete EDTA-free Protease Inhibitor. Clarified lysate was applied to Ni-NTA agarose beads (Qiagen), washed with buffer containing 25 mM HEPES (pH 7.5), 20 mM Imidazole (pH 7.5), 500 mM NaCl, 10% glycerol, 5 mM βME and eluted with 25 mM HEPES (pH 7.5), 400 mM Imidazole (pH 7.5), 1 M NaCl, 10% glycerol, 5 mM βME. Protein was further purified with size exclusion chromatography using a Superdex 200 column (Cytiva) in 25 mM HEPES (pH 7.5), 500 mM NaCl, 10% Glycerol and 1 mM DTT. Fractions with high purity were identified by SDS–PAGE and concentrated using a centrifugation filter with a 30 kDa cutoff (Amicon, Millipore). Absorbance at 280 nm was measured using the storage buffer as a blank, and protein concentration was calculated using the predicted extinction coefficient. Aliquots were flash-frozen in liquid nitrogen and stored at −80°C.

BL21(DE3) cells expressing his-sumo-FAK-deltaKinase were induced with 1 mM IPTG at 16°C for 18 hours. Cells were collected by centrifugation and lysed by cell disruption (Emulsiflex-C5, Avestin) in 20 mM Tris (pH 8.0), 20 mM imidazole, 300 mM NaCl, 5 mM βME, 0.1% NP-40, 10% glycerol, 1 mM PMSF, 1 μg/ml antipain, 1 μg/ml benzamidine, 1 μg/ml leupeptin, and 1 μg/ml pepstatin. Centrifugation-cleared lysate was applied to Ni-NTA agarose (Qiagen), washed with 20 mM Tris (pH 8.0), 20 mM imidazole, 300 mM NaCl, 5 mM βME, 0.1% NP-40, 10% glycerol, and eluted with 20 mM Tris (pH 8.0), 300 mM imidazole, 300 mM NaCl, 5 mM βME, 0.1% NP-40, 10% glycerol. Eluate was applied to a Source 15Q anion exchange column (Cytiva) and eluted with a gradient of 150 mM to 350 mM NaCl in 20 mM Tris (pH 8.0), 10% glycerol, 1 mM DTT. Fractions containing FAK were pooled, and we added 300 mM NaCl. The his-sumo tag was removed using Ulp1 protease treatment for 2 hours at room temperature. Then we concentrated the sample to 500 μL using Amicon Ultra Centrifugal Filter units (Millipore). Cleaved FAK-deltaKinase was purified by size exclusion chromatography using a Superdex 200 Increase column (Cytiva) in 50 mM Hepes (pH 7.5), 300 mM NaCl, 10% glycerol, 1mM DTT.

##### SARS-CoV-2 nucleocapsid protein

MBP-tagged full length SARS-CoV-2 nucleocapsid protein (expressed from Addgene plasmid #157867, a gift from Nicolas Fawzi) was expressed in *Escherichia coli* (*E. coli*) BL21(DE3) cells.

Recombinant protein was purified as described previously^82^, with minor modifications. Cells expressing SARS2-NCAP were harvested by centrifugation and resuspended in buffer containing 20 mM Tris-HCl (pH 8.0), 1 M NaCl, 10 mM imidazole, and cOmplete EDTA-free Protease Inhibitor (Roche). The samples were flash frozen and stored at −80°C. Cell suspension was thawed and lysed using cell disruption (Emulsiflex-C5, Avestin), and lysates were clarified by centrifugation at 51,428 *g* for 45 min at 4°C. The supernatant was filtered through a 0.2 µm filter and loaded onto a HisTrap affinity column equilibrated in buffer A (20 mM Tris-HCl pH 8.0, 1 M NaCl, 10 mM imidazole) using an ÄKTA system. Bound protein was eluted using buffer B containing 300 mM imidazole. Peak fractions were pooled and concentrated, followed by size-exclusion chromatography in SEC buffer consisting of 20 mM Tris-HCl (pH 8.0), 1 M NaCl, and 1 mM EDTA. Fractions with high purity were identified by SDS–PAGE and concentrated using a centrifugation filter with a 10 kDa cutoff (Amicon, Millipore). Absorbance at 280 nm was measured using the storage buffer as a blank, and protein concentration was calculated using the predicted extinction coefficient. Aliquots were flash-frozen in liquid nitrogen and stored at −80°C.

##### G3BP1

GFP-G3BP1 (gift from Parker Lab) protein was expressed in E coli Rosetta (DE3) cells. Recombinant protein was purified as described previously^83^, with minor modifications. Cells were harvested by centrifugation and resuspended in lysis buffer containing 50 mM HEPES (pH 7.5), 250 mM NaCl, and 1 mM DTT supplemented with cOmplete EDTA-free Protease Inhibitor (Roche). The samples were flash frozen and stored at −80°C. Cell suspension was thawed and lysed using cell disruption (Emulsiflex-C5, Avestin), and lysates were clarified by centrifugation at 51,428 *g* for 45 min at 4°C. The cleared lysate was loaded onto Glutathione Sepharose resin (Cytiva) equilibrated in lysis buffer. After washing, bound protein was eluted using lysis buffer supplemented with 10 mM reduced glutathione. The eluates were incubated with RNase A/T1 (Invitrogen) to reduce nucleic acid contamination. GST tags were removed by overnight TEV protease cleavage at room temperature. The cleaved protein was concentrated followed by size-exclusion chromatography on a Superdex 200 column equilibrated in SEC buffer containing 50 mM HEPES (pH 7.5), 400 mM NaCl, and 1 mM DTT. Fractions with high purity were identified by SDS–PAGE and concentrated using a centrifugation filter with a 30 kDa cutoff (Amicon, Millipore). Aliquots were snap-frozen in liquid nitrogen, and stored at −80°C.

##### TDP-43

TDP-43 was purified as described previously^84^. Briefly, His-MBP tagged full length TDP-43 (expressed from pJ4M/TDP-43; Addgene plasmid # 104480, a gift from Nicolas Fawzi) was transformed into One Shot BL21 Star (DE3) chemically competent *E. coli* (Thermo Fisher Scientific). Cells were then harvested by centrifugation for 20 min at 4,658 x *g* then flash frozen and stored at −80°C. Cell suspension was then thawed and resuspended in Buffer A (1 M NaCl, 20 mM Tris pH 8.0, 10 mM imidazole pH 8.0, 10% glycerol, 2.5 mM BME) supplemented with cOmplete EDTA-free Protease Inhibitor (Roche), followed by lysis via cell disruption (Emulsiflex-C5, Avestin). Lysates were then clarified by centrifugation at 51,428 *g* for 1 hr and supernatants filtered through a 0.2 µm filter. Lysates were then loaded onto a HisTrap column equilibrated with Buffer A using an ÄKTA chromatography system. Protein was eluted with buffer A containing 500 mM imidazole. Peak fractions were pooled and concentrated then loaded onto a SD200 26-600 column equilibrated with Buffer B (300 mM NaCl, 20 mM Tris pH 8.0, 1 mM DTT). Fractions with high purity were validated using SDS-PAGE, then pooled and concentrated with a 30 kDa cutoff Amicon filter. Samples were flash-frozen in liquid nitrogen and stored at −80°C.

## Supplementary Figures

**Figure S1.**
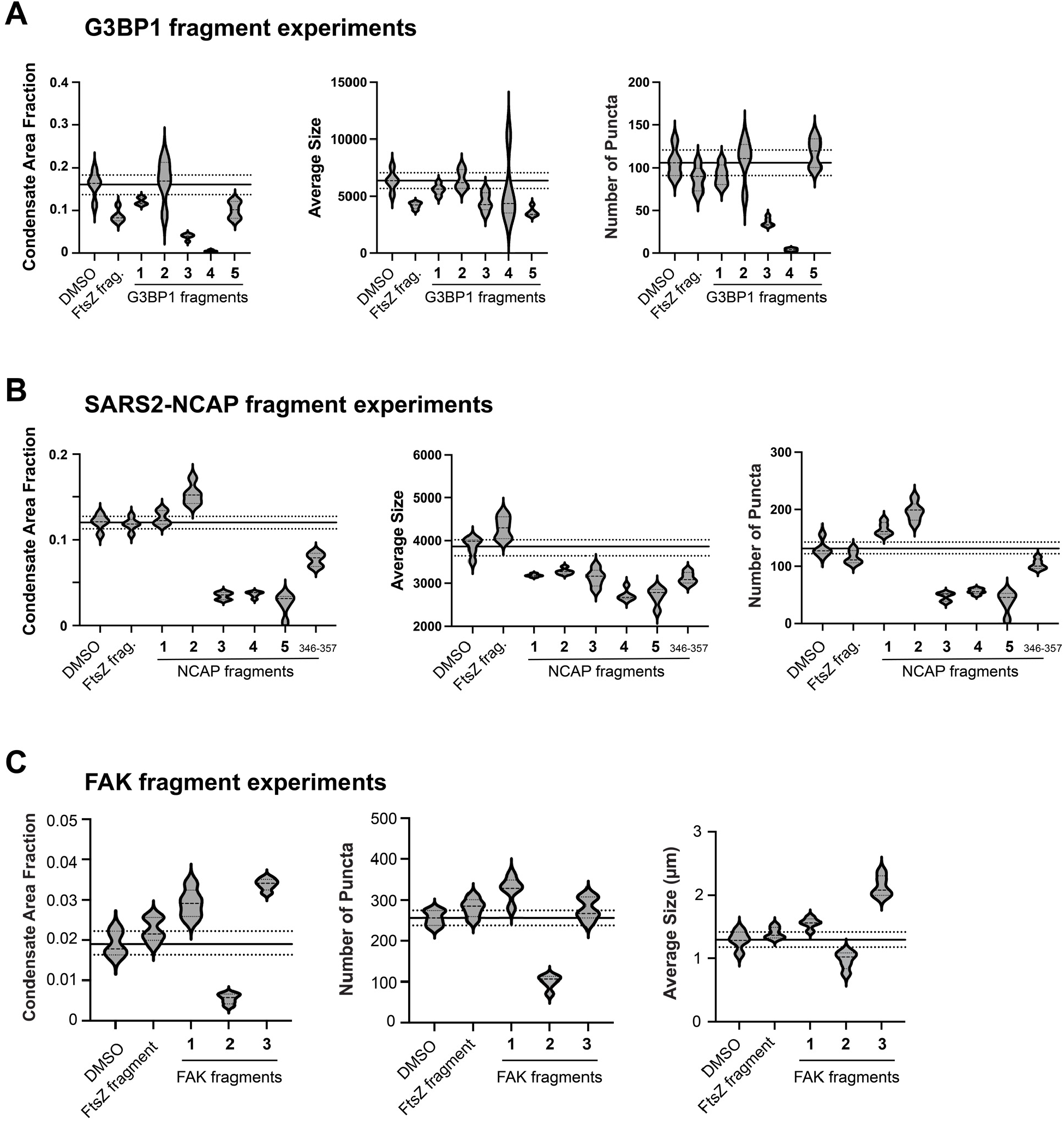
Additional quantifications of protein fragment effects on condensate formation. Measurements of protein fragment effects on condensate area fraction, average condensate size, and number of condensate puncta for G3BP1, SARS2-NCAP, and FAK, from experiments performed as described in **Fig. 2** and **Fig. 3**. Protein fragments are designated as in **Fig. 2** and **Fig. 3**. DMSO, DMSO control. FtsZ frag., experiments performed with an equal concentration of fragment 34-63 of the unrelated bacterial cytoskeletal protein FtsZ.

**Figure S2.**
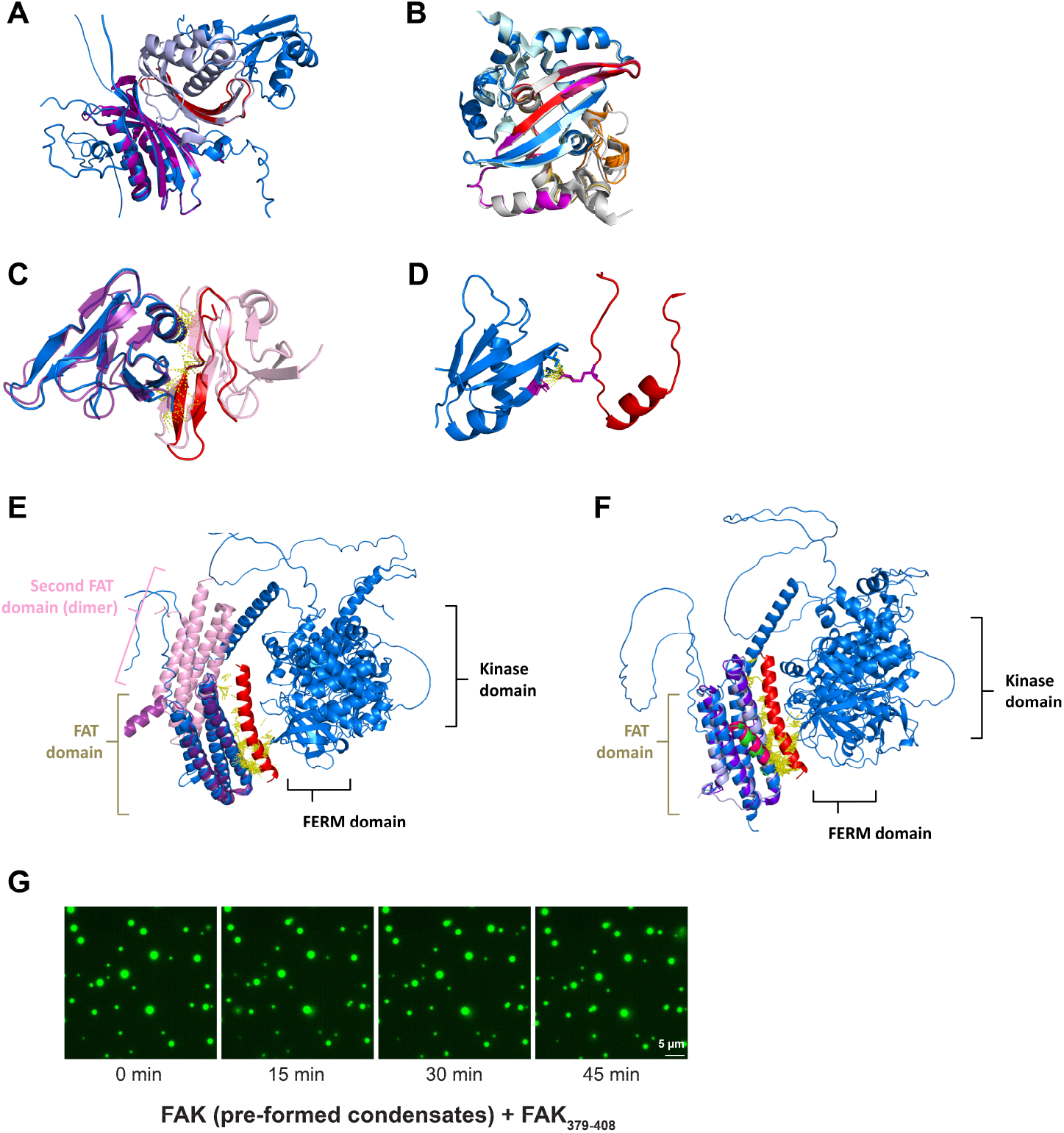
Additional FragFold model analysis reveals fragment binding at interactions driving condensate formation, and investigating effects of FAK fragments on pre-formed condensates. (**A**) G3BP1_86-115_ is predicted to form a native-like NTF2L dimer interaction. FragFold model of G3BP1, in blue, binding fragment G3BP1_86-115_ (red), overlaid with the experimental crystal structure of the G3BP1 NTF2L domain dimer (purple and light violet; PDB ID 3Q90). (**B**) SARS2-NCAP protein fragments are predicted to form native-like DBD dimer interactions. Overlay of the experimental crystal structure of the SARS2-NCAP dimerization domain (DD) (dimer in cyan and grey; PDB ID 6WZO) with FragFold models of the DD region NCAP, in blue, binding fragments NCAP_308-337_ (red), NCAP_257-286_ (orange), NCAP_290-319_ (gold), NCAP_304-333_ (magenta), and NCAP_328-357_ (pink). (**C – D**) Further details of TDP-43 fragment binding modes predicted with FragFold. (**C**) TDP-43_37-66_ is predicted to form a native-like dimerization interaction with the N-terminal oligomerization domain of TDP-43, and thereby blocking oligomerization. FragFold model of TDP-43, in blue, highlighting just the N-terminal domain, binding fragment TDP-43_37-66_ (red); overlaid with the experimentally determined structure of the TDP-43 N-terminal domain dimer, PDB ID 6B1G, with one monomer in purple, and the second in pink. (**D**) TDP-43_142-171_ is predicted to form an RRM1-RRM2 native-like interaction involving R151-D247. FragFold model of TDP-43, in blue, zooming in on the RRM2 domain, binding fragment TDP-43_142-171_ (red); residues R151 (of the fragment) and D247 (of the full-length protein) are highlighted in purple. (**E – F**) The FAK_977-1006_ binding mode predicted with FragFold is consistent with enhancement of FAT dimerization and of FAT-FERM interactions. (**E**) FAK fragment 977-1006 binds in a mode that would not interfere with FAT dimerization, and could enhance this interaction as a bridging fragment. FragFold model of focal adhesion kinase (FAK), in blue, binding fragment FAK_977-1006_ (red); overlaid with experimental structures of a FAK FAT domain dimer (one monomer purple, the other pink), PDB ID 5F28. (**F**) FAK fragment 977-1006 binds in a similar mode to paxillin LD2 and LD4. FragFold model of focal adhesion kinase (FAK), in blue, binding fragment FAK_977-1006_ (red); overlaid with experimental structures of FAK FAT domain (purple, light blue) bound to paxillin motifs LD2 (magenta, PDB ID 1OW8) and LD4 (green, PDB ID 3GM1). (**G**) Inhibitory FAK fragment 379-408 does not dissolve pre-formed FAK condensates *in vitro*. FAK condensates were pre-formed in the absence of protein fragments, at 1 µM FAK. Following this, fragment FAK_379-408_ was added at 10 µM after the indicated lag times.

**Figure S3.**
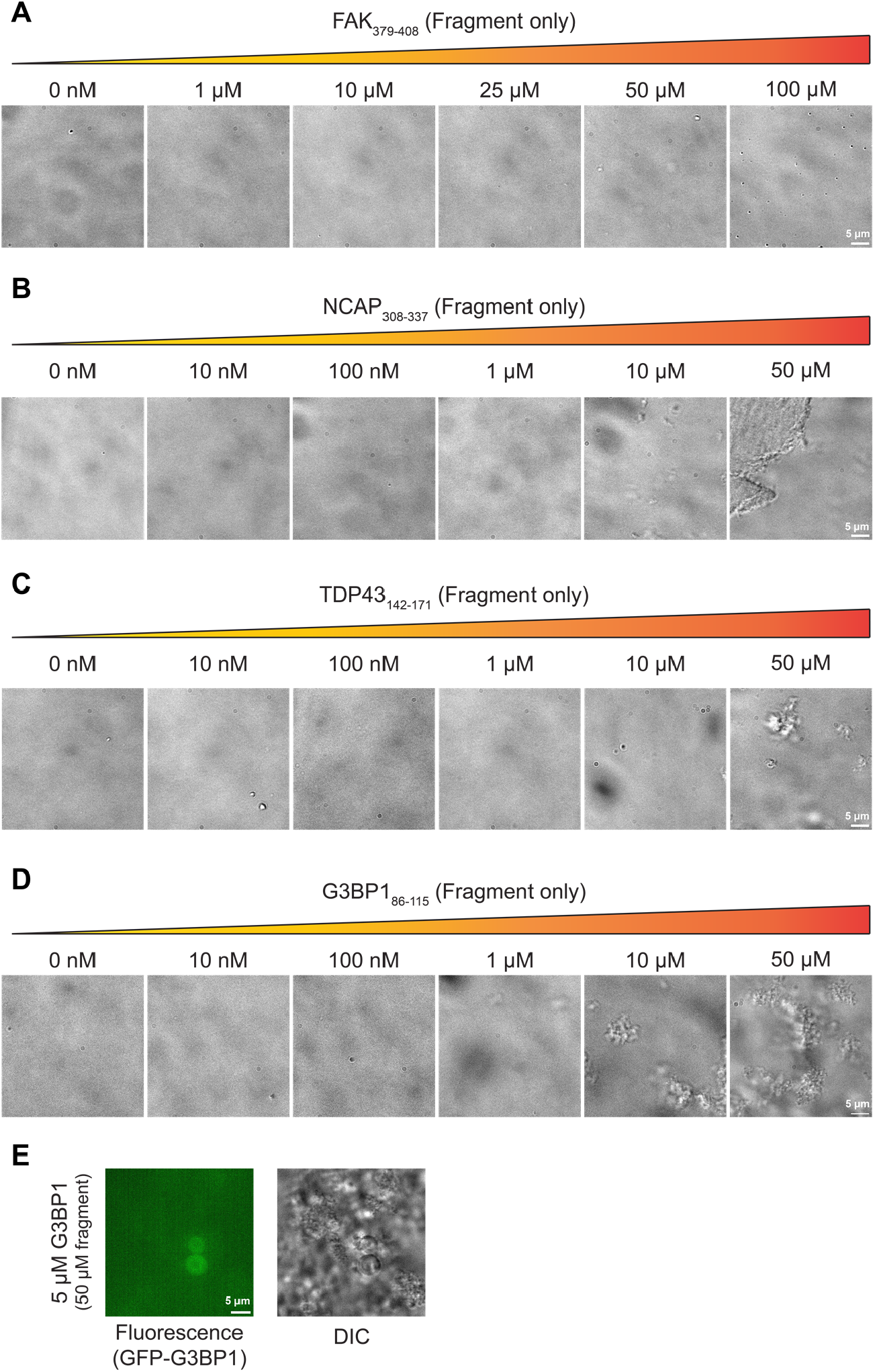
Aggregation propensities of protein fragments at high concentrations. Concentration series of inhibitory protein fragments of (**A**) FAK, (**B**) NCAP, (**C**) TDP-43, and (**D**) G3BP1 with no full-length protein present, imaged by DIC. (**E**) Inhibition of G3BP1 condensates by G3BP1 fragment 86-115, as in **Fig. 2B**, comparing view by epifluorescence (GFP-tagged G3BP1) and DIC imaging. G3BP1 is not enriched in fragment aggregates.

**Figure S4.**
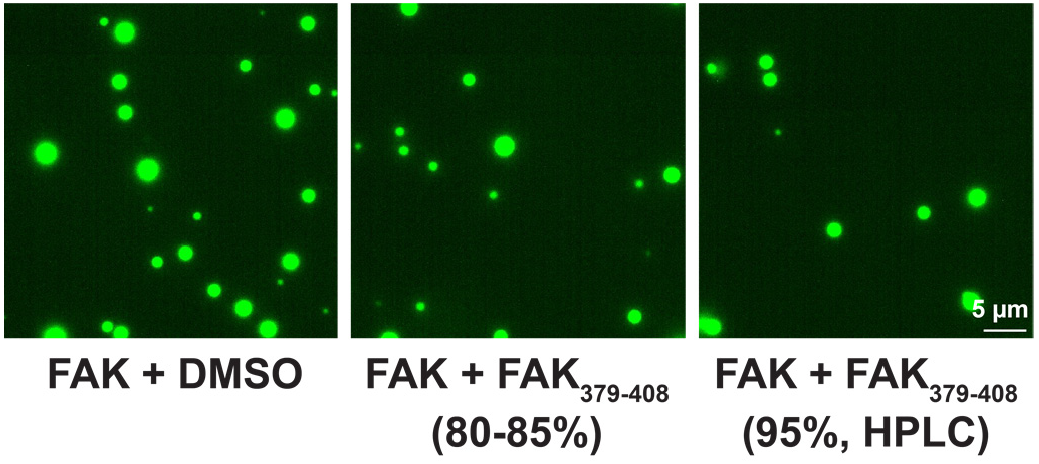
Inhibitory function of FAK_379-408_ is not reduced by further purification. *In vitro* condensate experiments with FAK and FAK_379-408_, performed as in **Fig. 2**, with either 80-85% purity fragments or further HPLC-purified fragments (95% purity). Both preparations inhibit FAK condensate formation.

**Table S1.**
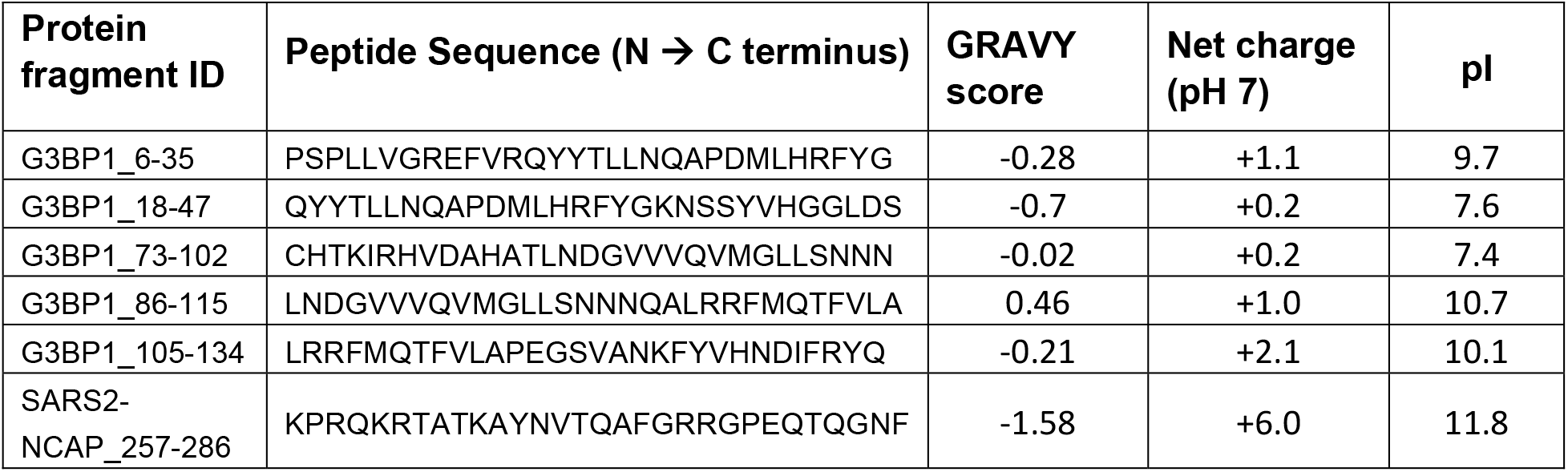
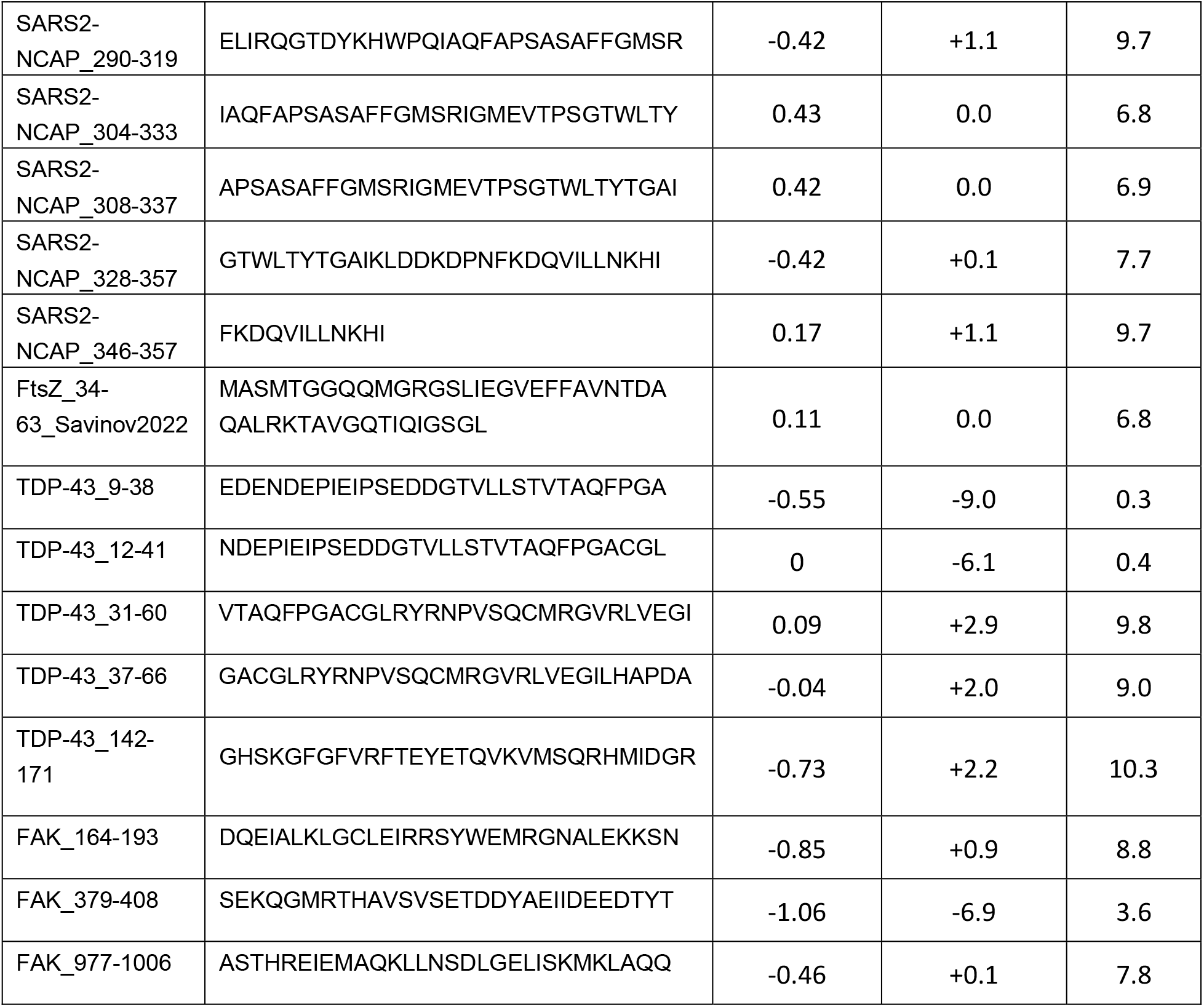
Sequences of protein fragments experimentally investigated in this work and their physicochemical properties. GRAVY (Grand Average of Hydropathy) scores were calculated using the Peptide Analyzing Tool from Thermo Fisher. Net charge at pH 7 and pI were calculated using the PepCalc tool from Innovagen.

## References

1. Feric, M., and Misteli, T. (2021). Phase separation in genome organization across evolution. Trends in Cell Biology 31, 671–685. 10.1016/j.tcb.2021.03.001.

2. Ladouceur, A.-M., Parmar, B.S., Biedzinski, S., Wall, J., Tope, S.G., Cohn, D., Kim, A., Soubry, N., Reyes-Lamothe, R., and Weber, S.C. (2020). Clusters of bacterial RNA polymerase are biomolecular condensates that assemble through liquid–liquid phase separation. Proceedings of the National Academy of Sciences 117, 18540–18549. 10.1073/pnas.2005019117.

3. Monterroso, B., Margolin, W., Boersma, A.J., Rivas, G., Poolman, B., and Zorrilla, S. (2024). Macromolecular Crowding, Phase Separation, and Homeostasis in the Orchestration of Bacterial Cellular Functions. Chem. Rev. 124, 1899–1949. 10.1021/acs.chemrev.3c00622.

4. Franzmann, T.M., and Alberti, S. (2019). Protein Phase Separation as a Stress Survival Strategy. Cold Spring Harb Perspect Biol 11, a034058. 10.1101/cshperspect.a034058.

5. Riback, J.A., Katanski, C.D., Kear-Scott, J.L., Pilipenko, E.V., Rojek, A.E., Sosnick, T.R., and Drummond, D.A. (2017). Stress-Triggered Phase Separation Is an Adaptive, Evolutionarily Tuned Response. Cell 168, 1028–1040.e19. 10.1016/j.cell.2017.02.027.

6. Zhu, S., Gu, J., Yao, J., Li, Y., Zhang, Z., Xia, W., Wang, Z., Gui, X., Li, L., Li, D., et al. (2022). Liquid-liquid phase separation of RBGD2/4 is required for heat stress resistance in Arabidopsis. Developmental Cell 57, 583–597.e6. 10.1016/j.devcel.2022.02.005.

7. Liu, X., Zhu, J.-K., and Zhao, C. (2023). Liquid-liquid phase separation as a major mechanism of plant abiotic stress sensing and responses. Stress Biology 3, 56. 10.1007/s44154-023-00141-x.

8. Jeon, S., Jeon, Y., Lim, J.-Y., Kim, Y., Cha, B., and Kim, W. (2025). Emerging regulatory mechanisms and functions of biomolecular condensates: implications for therapeutic targets. Sig Transduct Target Ther 10, 4. 10.1038/s41392-024-02070-1.

9. Xiao, Q., McAtee, C.K., and Su, X. (2022). Phase separation in immune signalling. Nat Rev Immunol 22, 188–199. 10.1038/s41577-021-00572-5.

10. Su, X., Ditlev, J.A., Hui, E., Xing, W., Banjade, S., Okrut, J., King, D.S., Taunton, J., Rosen, M.K., and Vale, R.D. (2016). Phase separation of signaling molecules promotes T cell receptor signal transduction. Science 352, 595–599. 10.1126/science.aad9964.

11. Banani, S.F., Lee, H.O., Hyman, A.A., and Rosen, M.K. (2017). Biomolecular condensates: organizers of cellular biochemistry. Nat Rev Mol Cell Biol 18, 285–298. 10.1038/nrm.2017.7.

12. Mittag, T., and Pappu, R.V. (2022). A conceptual framework for understanding phase separation and addressing open questions and challenges. Molecular Cell 82, 2201–2214. 10.1016/j.molcel.2022.05.018.

13. Borcherds, W., Bremer, A., Borgia, M.B., and Mittag, T. (2021). How do intrinsically disordered protein regions encode a driving force for liquid–liquid phase separation? Current Opinion in Structural Biology 67, 41–50. 10.1016/j.sbi.2020.09.004.

14. Ditlev, J.A., Case, L.B., and Rosen, M.K. (2018). Who’s In and Who’s Out—Compositional Control of Biomolecular Condensates. Journal of Molecular Biology 430, 4666–4684. 10.1016/j.jmb.2018.08.003.

15. Wang, B., Zhang, L., Dai, T., Qin, Z., Lu, H., Zhang, L., and Zhou, F. (2021). Liquid–liquid phase separation in human health and diseases. Sig Transduct Target Ther 6, 290. 10.1038/s41392-021-00678-1.

16. Alberti, S., and Dormann, D. (2019). Liquid–Liquid Phase Separation in Disease. Annual Review of Genetics 53, 171–194. 10.1146/annurev-genet-112618-043527.

17. Fu, Q., Zhang, B., Chen, X., and Chu, L. (2024). Liquid–liquid phase separation in Alzheimer’s disease. J Mol Med 102, 167–181. 10.1007/s00109-023-02407-3.

18. Mehta, S., and Zhang, J. (2022). Liquid–liquid phase separation drives cellular function and dysfunction in cancer. Nat Rev Cancer 22, 239–252. 10.1038/s41568-022-00444-7.

19. Tong, X., Tang, R., Xu, J., Wang, W., Zhao, Y., Yu, X., and Shi, S. (2022). Liquid–liquid phase separation in tumor biology. Sig Transduct Target Ther 7, 221. 10.1038/s41392-022-01076-x.

20. Cubuk, J., Alston, J.J., Incicco, J.J., Singh, S., Stuchell-Brereton, M.D., Ward, M.D., Zimmerman, M.I., Vithani, N., Griffith, D., Wagoner, J.A., et al. (2021). The SARS-CoV-2 nucleocapsid protein is dynamic, disordered, and phase separates with RNA. Nat Commun 12, 1936. 10.1038/s41467-021-21953-3.

21. Chau, B.-A., Chen, V., Cochrane, A.W., Parent, L.J., and Mouland, A.J. (2023). Liquid-liquid phase separation of nucleocapsid proteins during SARS-CoV-2 and HIV-1 replication. Cell Reports 42. 10.1016/j.celrep.2022.111968.

22. Li, P., Chen, P., Qi, F., Shi, J., Zhu, W., Li, J., Zhang, P., Xie, H., Li, L., Lei, M., et al. (2024). High-throughput and proteome-wide discovery of endogenous biomolecular condensates. Nat. Chem. 16, 1101–1112. 10.1038/s41557-024-01485-1.

23. Norrild, R.K., Bülow, S. von, Halldórsson, E., Lindorff-Larsen, K., Rogers, J.M., and Buell, A.K. (2025). Proteome-scale quantification of the interactions driving condensate formation of intrinsically disordered proteins. Preprint at bioRxiv, 10.1101/2024.12.21.629870 https://doi.org/10.1101/2024.12.21.629870.

24. Kappel, K., Strebinger, D., Edmonds, K.K., Chau-Duy-Tam VO, S., Vockley, C.M., Biswas, T., Farhi, S.L., Macrae, R., Zhang, F., and Regev, A. (2025). Characterizing protein sequence determinants of nuclear condensates by high-throughput pooled imaging with CondenSeq. Nat Methods 22, 1464–1475. 10.1038/s41592-025-02726-y.

25. von Bülow, S., Tesei, G., Zaidi, F.K., Mittag, T., and Lindorff-Larsen, K. (2025). Prediction of phase-separation propensities of disordered proteins from sequence. Proceedings of the National Academy of Sciences 122, e2417920122. 10.1073/pnas.2417920122.

26. He, Y., Ongwae, G.M., Mondal, A., Moses, J.A., Mittal, J., and Pires, M.M. (2026). A high-throughput, flow cytometry approach to measure phase behavior and exchange in biomolecular condensates. Nat Commun 17, 1337. 10.1038/s41467-025-68093-6.

27. Brangwynne, C.P., Eckmann, C.R., Courson, D.S., Rybarska, A., Hoege, C., Gharakhani, J., Jülicher, F., and Hyman, A.A. (2009). Germline P Granules Are Liquid Droplets That Localize by Controlled Dissolution/Condensation. Science 324, 1729–1732. 10.1126/science.1172046.

28. Li, P., Banjade, S., Cheng, H.-C., Kim, S., Chen, B., Guo, L., Llaguno, M., Hollingsworth, J.V., King, D.S., Banani, S.F., et al. (2012). Phase transitions in the assembly of multivalent signalling proteins. Nature 483, 336–340. 10.1038/nature10879.

29. Molliex, A., Temirov, J., Lee, J., Coughlin, M., Kanagaraj, A.P., Kim, H.J., Mittag, T., and Taylor, J.P. (2015). Phase Separation by Low Complexity Domains Promotes Stress Granule Assembly and Drives Pathological Fibrillization. Cell 163, 123–133. 10.1016/j.cell.2015.09.015.

30. Das, R.K., Ruff, K.M., and Pappu, R.V. (2015). Relating sequence encoded information to form and function of intrinsically disordered proteins. Current Opinion in Structural Biology 32, 102–112. 10.1016/j.sbi.2015.03.008.

31. Farag, M., Borcherds, W.M., Bremer, A., Mittag, T., and Pappu, R.V. (2023). Phase separation of protein mixtures is driven by the interplay of homotypic and heterotypic interactions. Nat Commun 14, 5527. 10.1038/s41467-023-41274-x.

32. Liu, C., Wu, K., Choi, H., Han, H.L., Zhang, X., Watson, J.L., Ahn, G., Zhang, J.Z., Shijo, S., Good, L.L., et al. (2025). Diffusing protein binders to intrinsically disordered proteins. Nature 644, 809–817. 10.1038/s41586-025-09248-9.

33. Stark, H., Faltings, F., Choi, M., Xie, Y., Hur, E., O’Donnell, T., Bushuiev, A., Uçar, T., Passaro, S., Mao, W., et al. (2025). BoltzGen: Toward Universal Binder Design. Preprint at bioRxiv, 10.1101/2025.11.20.689494 https://doi.org/10.1101/2025.11.20.689494.

34. Harmon, T.S., Holehouse, A.S., and Pappu, R.V. (2018). Differential solvation of intrinsically disordered linkers drives the formation of spatially organized droplets in ternary systems of linear multivalent proteins. New J. Phys. 20, 045002. 10.1088/1367-2630/aab8d9.

35. Ruff, K.M., Dar, F., and Pappu, R.V. (2021). Ligand effects on phase separation of multivalent macromolecules. Proceedings of the National Academy of Sciences 118, e2017184118. 10.1073/pnas.2017184118.

36. Martin, E.W., and Mittag, T. (2018). Relationship of Sequence and Phase Separation in Protein Low-Complexity Regions. Biochemistry 57, 2478–2487. 10.1021/acs.biochem.8b00008.

37. von Hofe, J., Abacousnac, J., Chen, M., Sasazawa, M., Javér Kristiansen, I., Westrey, S., Grier, D.G., and Saurabh, S. (2025). Multivalency Controls the Growth and Dynamics of a Biomolecular Condensate. J. Am. Chem. Soc. 147, 25242–25253. 10.1021/jacs.5c02947.

38. Anzai, R., Mabuchi, A., and Hata, S. Coiled-coils as emerging drivers of liquid–liquid phase separation.

39. Poudyal, M., Patel, K., Gadhe, L., Sawner, A.S., Kadu, P., Datta, D., Mukherjee, S., Ray, S., Navalkar, A., Maiti, S., et al. (2023). Intermolecular interactions underlie protein/peptide phase separation irrespective of sequence and structure at crowded milieu. Nat Commun 14, 6199. 10.1038/s41467-023-41864-9.

40. Nie, J., Zhang, X., Hu, Z., Wang, W., Schroer, M.A., Ren, J., Svergun, D., Chen, A., Yang, P., and Zeng, A.-P. (2025). A globular protein exhibits rare phase behavior and forms chemically regulated orthogonal condensates in cells. Nat Commun 16, 2449. 10.1038/s41467-025-57886-4.

41. Savinov, A., Swanson, S., Keating, A.E., and Li, G.-W. (2025). High-throughput discovery of inhibitory protein fragments with AlphaFold. Proceedings of the National Academy of Sciences 122, e2322412122. 10.1073/pnas.2322412122.

42. Savinov, A., Fernandez, A., and Fields, S. (2022). Mapping functional regions of essential bacterial proteins with dominant-negative protein fragments. Proceedings of the National Academy of Sciences 119, e2200124119. 10.1073/pnas.2200124119.

43. Wang, S., Dai, T., Qin, Z., Pan, T., Chu, F., Lou, L., Zhang, L., Yang, B., Huang, H., Lu, H., et al. (2021). Targeting liquid–liquid phase separation of SARS-CoV-2 nucleocapsid protein promotes innate antiviral immunity by elevating MAVS activity. Nat Cell Biol 23, 718–732. 10.1038/s41556-021-00710-0.

44. Dorrity, M.W., Queitsch, C., and Fields, S. (2019). High-throughput identification of dominant negative polypeptides in yeast. Nat Methods 16, 413–416. 10.1038/s41592-019-0368-0.

45. Savinov, A., and Roth, F.P. (2021). Seeds of their own destruction: Dominant-negative peptide screening yields functional insight and therapeutic leads. Cell Systems 12, 691–693. 10.1016/j.cels.2021.06.003.

46. Ford, K.M., Panwala, R., Chen, D.-H., Portell, A., Palmer, N., and Mali, P. (2021). Peptide-tiling screens of cancer drivers reveal oncogenic protein domains and associated peptide inhibitors. Cell Systems 12, 716–732.e7. 10.1016/j.cels.2021.05.002.

47. Ge, Y., Jin, J., Li, J., Ye, M., and Jin, X. (2022). The roles of G3BP1 in human diseases (review). Gene 821, 146294. 10.1016/j.gene.2022.146294.

48. Matsuki, H., Takahashi, M., Higuchi, M., Makokha, G.N., Oie, M., and Fujii, M. (2013). Both G3BP1 and G3BP2 contribute to stress granule formation. Genes to Cells 18, 135–146. 10.1111/gtc.12023.

49. Wu, W., Cheng, Y., Zhou, H., Sun, C., and Zhang, S. (2023). The SARS-CoV-2 nucleocapsid protein: its role in the viral life cycle, structure and functions, and use as a potential target in the development of vaccines and diagnostics. Virol J 20, 6. 10.1186/s12985-023-01968-6.

50. Sun, Y., and Chakrabartty, A. (2017). Phase to Phase with TDP-43. Biochemistry 56, 809–823. 10.1021/acs.biochem.6b01088.

51. Meneses, A., Koga, S., O’Leary, J., Dickson, D.W., Bu, G., and Zhao, N. (2021). TDP-43 Pathology in Alzheimer’s Disease. Mol Neurodegeneration 16, 84. 10.1186/s13024-021-00503-x.

52. Boer, E.M.J. de, Orie, V.K., Williams, T., Baker, M.R., Oliveira, H.M.D., Polvikoski, T., Silsby, M., Menon, P., Bos, M. van den, Halliday, G.M., et al. (2021). TDP-43 proteinopathies: a new wave of neurodegenerative diseases. J Neurol Neurosurg Psychiatry 92, 86–95. 10.1136/jnnp-2020-322983.

53. Vognsen, T., Møller, I.R., and Kristensen, O. (2013). Crystal Structures of the Human G3BP1 NTF2-Like Domain Visualize FxFG Nup Repeat Specificity. PLOS ONE 8, e80947. 10.1371/journal.pone.0080947.

54. Yang, P., Mathieu, C., Kolaitis, R.-M., Zhang, P., Messing, J., Yurtsever, U., Yang, Z., Wu, J., Li, Y., Pan, Q., et al. (2020). G3BP1 Is a Tunable Switch that Triggers Phase Separation to Assemble Stress Granules. Cell 181, 325–345.e28. 10.1016/j.cell.2020.03.046.

55. Schulte, T., Liu, L., Panas, M.D., Thaa, B., Dickson, N., Götte, B., Achour, A., and McInerney, G.M. (2016). Combined structural, biochemical and cellular evidence demonstrates that both FGDF motifs in alphavirus nsP3 are required for efficient replication. Open Biol 6, 160078. 10.1098/rsob.160078.

56. Somasekharan, S.P., and Gleave, M. (2021). SARS-CoV-2 nucleocapsid protein interacts with immunoregulators and stress granules and phase separates to form liquid droplets. FEBS Letters 595, 2872–2896. 10.1002/1873-3468.14229.

57. Ye, Q., West, A.M.V., Silletti, S., and Corbett, K.D. (2020). Architecture and self-assembly of the SARS-CoV-2 nucleocapsid protein. Protein Science 29, 1890–1901. 10.1002/pro.3909.

58. Carter, G.C., Hsiung, C.-H., Simpson, L., Yang, H., and Zhang, X. (2021). N-terminal Domain of TDP43 Enhances Liquid-Liquid Phase Separation of Globular Proteins. Journal of Molecular Biology 433, 166948. 10.1016/j.jmb.2021.166948.

59. Zacco, E., Graña-Montes, R., Martin, S.R., de Groot, N.S., Alfano, C., Tartaglia, G.G., and Pastore, A. (2019). RNA as a key factor in driving or preventing self-assembly of the TAR DNA-binding protein 43. Journal of Molecular Biology 431, 1671–1688. 10.1016/j.jmb.2019.01.028.

60. Huang, Y.-C., Lin, K.-F., He, R.-Y., Tu, P.-H., Koubek, J., Hsu, Y.-C., and Huang, J.J.-T. (2013). Inhibition of TDP-43 Aggregation by Nucleic Acid Binding. PLOS ONE 8, e64002. 10.1371/journal.pone.0064002.

61. Liu, W., Li, C., Shan, J., Wang, Y., and Chen, G. (2021). Insights into the aggregation mechanism of RNA recognition motif domains in TDP-43: a theoretical exploration. R Soc Open Sci. 8, 210160. 10.1098/rsos.210160.

62. Zhao, X., and Guan, J.-L. (2011). Focal adhesion kinase and its signaling pathways in cell migration and angiogenesis. Adv Drug Deliv Rev 63, 610–615. 10.1016/j.addr.2010.11.001.

63. Schlaepfer, D.D., and Mitra, S.K. (2004). Multiple connections link FAK to cell motility and invasion. Current Opinion in Genetics & Development 14, 92–101. 10.1016/j.gde.2003.12.002.

64. Schlaepfer, D.D., Ojalill, M., and Stupack, D.G. (2024). Focal adhesion kinase signaling – tumor vulnerabilities and clinical opportunities. J Cell Sci 137, jcs261723. 10.1242/jcs.261723.

65. Murphy, J.M., Rodriguez, Y.A.R., Jeong, K., Ahn, E.-Y.E., and Lim, S.-T.S. (2020). Targeting focal adhesion kinase in cancer cells and the tumor microenvironment. Exp Mol Med 52, 877–886. 10.1038/s12276-020-0447-4.

66. Sulzmaier, F.J., Jean, C., and Schlaepfer, D.D. (2014). FAK in cancer: mechanistic findings and clinical applications. Nat Rev Cancer 14, 598–610. 10.1038/nrc3792.

67. Haanen, T.J., and Schlaepfer, D.D. (2026). Disarming cancer resistance: FAK as a therapeutic target. Trends in Cancer. 10.1016/j.trecan.2025.12.008.

68. Case, L.B., De Pasquale, M., Henry, L., and Rosen, M.K. Synergistic phase separation of two pathways promotes integrin clustering and nascent adhesion formation. eLife 11, e72588. 10.7554/eLife.72588.

69. Kumar, A., Tanaka, K., and Schwartz, M.A. (2025). Focal adhesion-derived liquid-liquid phase separations regulate mRNA translation. eLife 13, RP96157. 10.7554/eLife.96157.

70. Colombo, G., Salem, I., Szczepski, K., Yu, P., Alfaiyz, S., Guzmán-Vega, F.J., Abogosh, A., Kulmanov, M., Al-Harthi, S., Kadaré, G., et al. (2026). Molecular basis and cellular effects of Janus-class–driven cytoplasmic PYK2 coacervates. Commun Biol 9, 186. 10.1038/s42003-025-09463-0.

71. Lea, N.E., and Case, L.B. (2026). Kinase condensates enrich ATP and trigger autophosphorylation. Cell Reports 117459, in press (2026).

72. Cardoso, A.C., Pereira, A.H.M., Ambrosio, A.L.B., Consonni, S.R., Rocha de Oliveira, R., Bajgelman, M.C., Dias, S.M.G., and Franchini, K.G. (2016). FAK Forms a Complex with MEF2 to Couple Biomechanical Signaling to Transcription in Cardiomyocytes. Structure 24, 1301–1310. 10.1016/j.str.2016.06.003.

73. Brami-Cherrier, K., Gervasi, N., Arsenieva, D., Walkiewicz, K., Boutterin, M., Ortega, A., Leonard, P.G., Seantier, B., Gasmi, L., Bouceba, T., et al. (2014). FAK dimerization controls its kinase-dependent functions at focal adhesions. EMBO J 33, 356–370. 10.1002/embj.201386399.

74. Lietha, D., Cai, X., Ceccarelli, D.F.J., Li, Y., Schaller, M.D., and Eck, M.J. (2007). Structural Basis for the Autoinhibition of Focal Adhesion Kinase. Cell 129, 1177–1187. 10.1016/j.cell.2007.05.041.

75. Kim, J.H., Lee, S.-R., Li, L.-H., Park, H.-J., Park, J.-H., Lee, K.Y., Kim, M.-K., Shin, B.A., and Choi, S.-Y. (2011). High Cleavage Efficiency of a 2A Peptide Derived from Porcine Teschovirus-1 in Human Cell Lines, Zebrafish and Mice. PLOS ONE 6, e18556. 10.1371/journal.pone.0018556.

76. Li, X., Zuber, P.K., Loughran, G., Bhatt, P.R., Alquraish, F., Ramakrishnan, V., Firth, A.E., and Atkins, J.F. (2026). Molecular architecture and diversity of StopGo/2A translational recoding. Proceedings of the National Academy of Sciences 123, e2528667123. 10.1073/pnas.2528667123.

77. Guillén-Boixet, J., Kopach, A., Holehouse, A.S., Wittmann, S., Jahnel, M., Schlüßler, R., Kim, K., Trussina, I.R.E.A., Wang, J., Mateju, D., et al. (2020). RNA-Induced Conformational Switching and Clustering of G3BP Drive Stress Granule Assembly by Condensation. Cell 181, 346–361.e17. 10.1016/j.cell.2020.03.049.

78. Babinchak, W.M., Haider, R., Dumm, B.K., Sarkar, P., Surewicz, K., Choi, J.-K., and Surewicz, W.K. (2019). The role of liquid–liquid phase separation in aggregation of the TDP-43 low-complexity domain. J Biol Chem 294, 6306–6317. 10.1074/jbc.RA118.007222.

79. Eschbach, E., Deibler, K., Korani, D., and Swanson, S. (2025). Predicting peptide aggregation with protein language model embeddings. Preprint at bioRxiv, 10.1101/2025.09.26.678773 http://doi.org/10.1101/2025.09.26.678773.

80. Stringer, C., Wang, T., Michaelos, M., and Pachitariu, M. (2021). Cellpose: a generalist algorithm for cellular segmentation. Nat Methods 18, 100–106. 10.1038/s41592-020-01018-x.

81. Pachitariu, M., and Stringer, C. (2022). Cellpose 2.0: how to train your own model. Nat Methods 19, 1634–1641. 10.1038/s41592-022-01663-4.

82. Perdikari, T.M., Murthy, A.C., Ryan, V.H., Watters, S., Naik, M.T., and Fawzi, N.L. (2020). SARS-CoV-2 nucleocapsid protein phase-separates with RNA and with human hnRNPs. EMBO J 39, EMBJ2020106478. 10.15252/embj.2020106478.

83. Parker, D.M., Tauber, D., and Parker, R. (2025). G3BP1 promotes intermolecular RNA-RNA interactions during RNA condensation. Molecular Cell 85, 571–584.e7. 10.1016/j.molcel.2024.11.012.

84. Hallegger, M., Chakrabarti, A.M., Lee, F.C.Y., Lee, B.L., Amalietti, A.G., Odeh, H.M., Copley, K.E., Rubien, J.D., Portz, B., Kuret, K., et al. (2021). TDP-43 condensation properties specify its RNA-binding and regulatory repertoire. Cell 184, 4680–4696.e22. 10.1016/j.cell.2021.07.018.

